# Gene Knock Up via 3’UTR editing to study gene function *in vivo*

**DOI:** 10.1101/775031

**Authors:** Kärt Mätlik, Soophie Olfat, Daniel R. Garton, Ana Montaño-Rodriguez, Giorgio Turconi, L. Lauriina Porokuokka, Anne Panhelainen, Nadine Schweizer, Jaakko Kopra, Mark C. Cowlishaw, T. Petteri Piepponen, Fu-Ping Zhang, Petra Sipilä, Johan Jakobsson, Jaan-Olle Andressoo

## Abstract

Currently available genetic tools do not allow researchers to upregulate (‘Knock Up’) the levels of a given protein while retaining its cell-type-specific regulation. As a result, we have limited ability to develop overexpression-related disease models, to study the contribution of single genes in diseases caused by copy number variations and to identify disease pathways for drug targets. Here we develop two approaches for endogenous gene upregulation: <u>c</u>onditional <u>K</u>nock <u>U</u>p (cKU) utilizing the Cre/lox system, and CRISPR-Cas9 mediated gene <u>K</u>nock <u>U</u>p (KU) in wild-type mouse embryos and human cells. Using glial cell line derived neurotrophic factor (GDNF) as a proof of concept, we show that both approaches resulted in upregulation of endogenous GDNF levels without disturbing *Gdnf*’s expression pattern. Furthermore, CNS-specific GDNF cKU resulted in dopaminergic abnormalities and schizophrenia-like phenotypes. Our results suggest that gene Knock Up can reveal unknown gene functions and provide novel entry points for studying neurological disease.

## Background

Understanding how aberrations in gene functioning lead to diseases and how the activity of endogenous signalling pathways can be harnessed to lead to different phenotypic outcomes and therapies is one of the major challenges for biomedical research and drug industry. One of the obstacles along this path is our limited ability to upregulate specific genes in endogenous signalling pathways—either by genetic tools or pharmacological compounds—to study gene function and to identify the genes or pathways that are beneficial if upregulated in a disease. Knocking out a gene or reducing the activity of the encoded protein is relatively achievable with various genetic tools. However, increasing the expression of specific genes while maintaining their natural expression pattern is much more of a challenge. Although transgenic animals and viral overexpression methods are widely used for this purpose, they usually result in gene expression levels that far exceed that of endogenous genes, and importantly, transgene expression patterns do not usually follow the expression of native genes^1^. Yet, human studies have shown that even as little as a 50-100% increase in endogenous gene expression levels or activity is sufficient to cause neurological disease^2–5^. Importantly, many of the genes implicated in these diseases encode regulatory proteins with restricted spatiotemporal expression patterns, further limiting the use of transgenic overexpression tools. Therefore, a tool that would allow researchers to conditionally increase gene expression levels within a physiological range, while retaining the correct expression pattern and transcriptional regulation, would be useful in many fields of basic and biomedical research, as well as for drug development.

With this in mind, we reasoned that in order to maintain correct gene expression patterns, modifications should be targeted towards regions that regulate expression post-transcriptionally, i.e., at the level of mRNA or protein. We focused on the 3’ untranslated region (3’UTR), because this region is well known for its role in regulating mRNA stability^6^. We hypothesised that by conditionally replacing negative regulation elements residing in the 3’UTR we could ‘Knock Up’ endogenous gene expression levels without altering the gene’s spatiotemporal expression pattern. For a proof of concept, we generated a <u>c</u>onditional <u>K</u>nock <u>U</u>p (cKU) model for the neurotrophic factor GDNF (glial cell line-derived neurotrophic factor), whose levels we have previously shown to be negatively regulated through its 3’UTR *in vitro* and *in vivo*^7^. GDNF is critically important for the development of peripheral tissues, including the kidney^8–11^. In addition, GDNF is a potent neurotrophic factor for midbrain dopamine neurons in culture^12^ and promotes dopamine synthesis and dopaminergic neuron fibre outgrowth if applied ectopically into the brain^13^. Increased dopamine signalling is associated with different neuropsychiatric diseases, including schizophrenia, where increased levels of presynaptic dopamine in the striatum are commonly observed^14–16^. Therefore, we were particularly interested in the outcome of brain-specific elevation of endogenous GDNF levels on brain dopamine system function, and the potential relevance of altered dopamine signalling in the context of schizophrenia.

To be able to specifically increase GDNF expression in the brain, we utilized the FLEx cassette^17^ to conditionally replace the Gdnf 3’UTR with a 3’UTR and polyadenylation signal derived from the bovine growth hormone (bGHpA), using Cre-mediated recombination. We found increased GDNF expression in GDNF cKU mice, while *Gdnf* expression sites appeared unchanged. Mice with endogenous GDNF upregulation in the central nervous system (CNS) exhibited increased presynaptic dopamine signalling in the striatum, and reduced dopamine levels and altered gene expression in prefrontal cortex, resembling findings from schizophrenic patients and animal models^14,15,18,19^. Interestingly, we found that reduction in prefrontal cortex dopamine levels can also be triggered by striatal upregulation of endogenous GDNF in adult mice using striatal AAV-Cre delivery. Finally, we asked if endogenous gene expression levels can be elevated in wild-type cells by 3’UTR editing. We found that excising inhibitory regions from the Gdnf 3’UTR using the CRISPR-Cas9 system increased endogenous GDNF expression levels in wild-type mouse embryos and in human cells. Taken together, our results suggest that gene Knock Up via 3’UTR editing is applicable to human cells and wild-type mice and can reveal novel information about gene function.

## Results

### Generation of <u>c</u>onditional <u>K</u>nock <u>U</u>p (cKU) allele

Gene overexpression levels and pattern in transgenic animals are mainly dependent upon transgene copy number, specificity of used promoter, integration site, and epigenetic changes across generations. As a result, the extent of overexpression and expression pattern of transgenes usually differ significantly from those of endogenous gene products^1^. Therefore, it can be difficult to draw conclusions about the functions of proteins, whose levels and expression pattern are tightly controlled. We hypothesised that by conditionally preventing post-transcriptional negative regulation of mRNA levels by 3’UTR replacement, endogenous mRNA and protein levels could be increased without changing the gene’s spatiotemporal expression pattern (Fig. 1a). We further hypothesised that by utilizing the Cre/lox system, an increase in endogenous gene levels could also be restricted to specific tissues or cell types. Notably, by only affecting post-transcriptional regulation, we expected that gene expression would not be induced in cells that do not normally express the gene (Fig. 1a).

**Figure 1.**
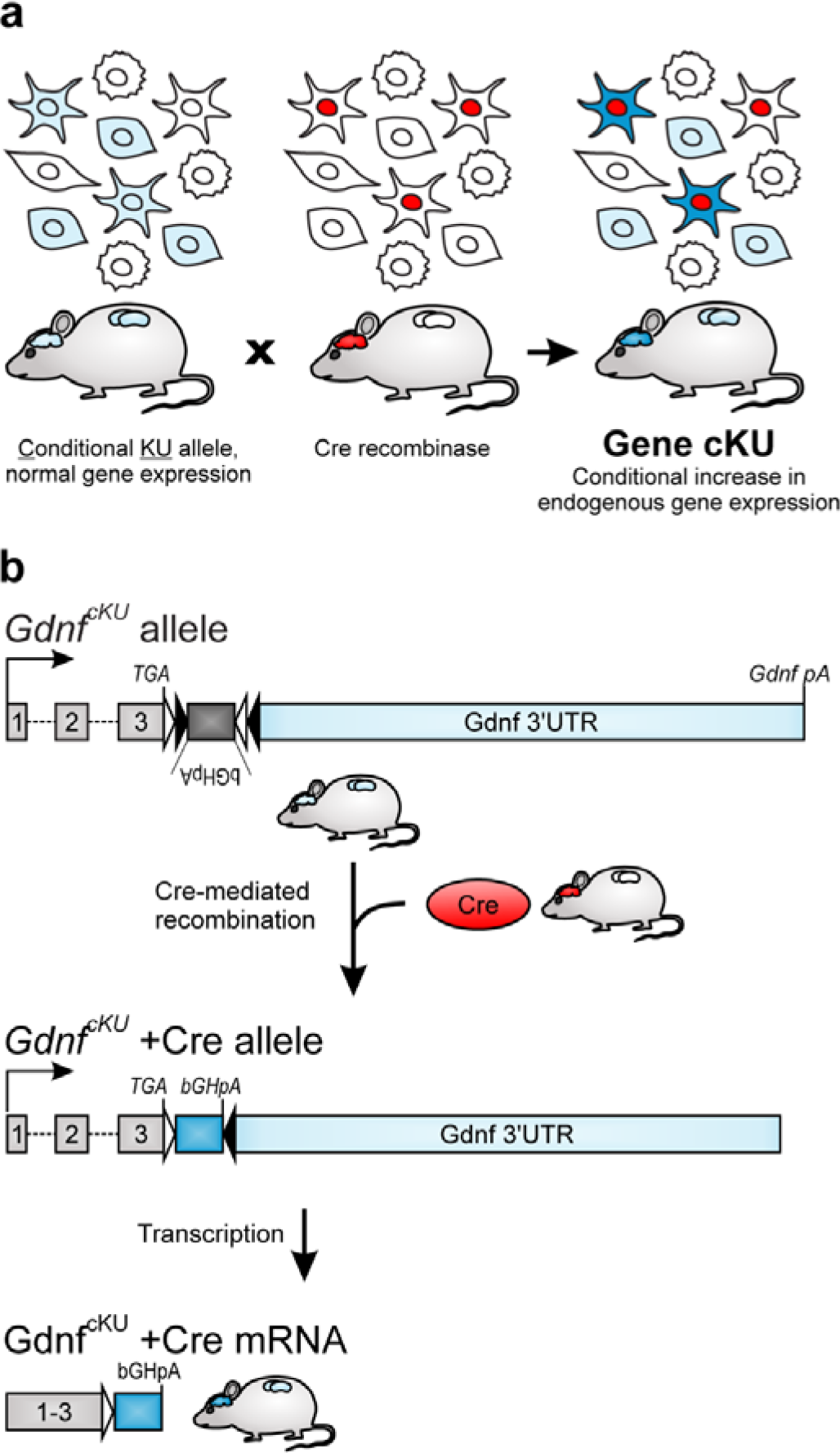
Schematic of <u>c</u>onditional <u>K</u>nock <u>U</u>p (cKU) approach. **a** Principle of conditional Knock Up of endogenous gene expression in natively expressing cells using the cKU allele combined with the Cre/lox system. **b** Schematic of *Gdnf*^*cKU*^ allele (top panel) and the resulting mRNA carrying a short 3’UTR (bottom panel) devoid of miR binding sites.

To provide a proof of concept for the approach, we generated a <u>c</u>onditional <u>K</u>nock <u>U</u>p (cKU) allele for glial cell line-derived neurotrophic factor (GDNF) by inserting a FLEx cassette^17^ containing bGHpA—a 3’UTR sequence previously shown to lead to high expression from the cognate mRNA^20^—in an inverted orientation immediately downstream of the stop codon of the *Gdnf* gene (Fig. 1b, Extended Data Fig. 1a). To evaluate the extent and expression pattern of conditional GDNF overexpression in *Gdnf*^*cKU*^ mice, we first analysed the kidneys, where *Gdnf*’s expression pattern and the outcomes of *Gdnf* gene deletion and increased levels of endogenous GDNF have been well characterized. Mice lacking GDNF have no kidneys^8–11^, whereas endogenous GDNF overexpression in GDNF hypermorphic mice causes a severe reduction in kidney size due to reduced migration of progenitors from the ureteric bud tip to the ureteric bud trunk^21^, and other histological alterations^7^. We first crossed *Gdnf*^*cKU*^ mice to a Pgk1-Cre mouse line, where Cre expression results in an early and uniform recombination^22^. We found that Gdnf mRNA and GDNF protein levels were significantly increased in the kidneys of *Gdnf*^*WT/cKU*^;Pgk1-Cre mice compared to wild-type mice at postnatal day (P) 3 (Fig. 2a-b). Similar to what we previously observed in GDNF hypermorphs^7^, kidney size was severely reduced in mice expressing increased levels of GDNF (Fig. 2c).

**Figure 2.**
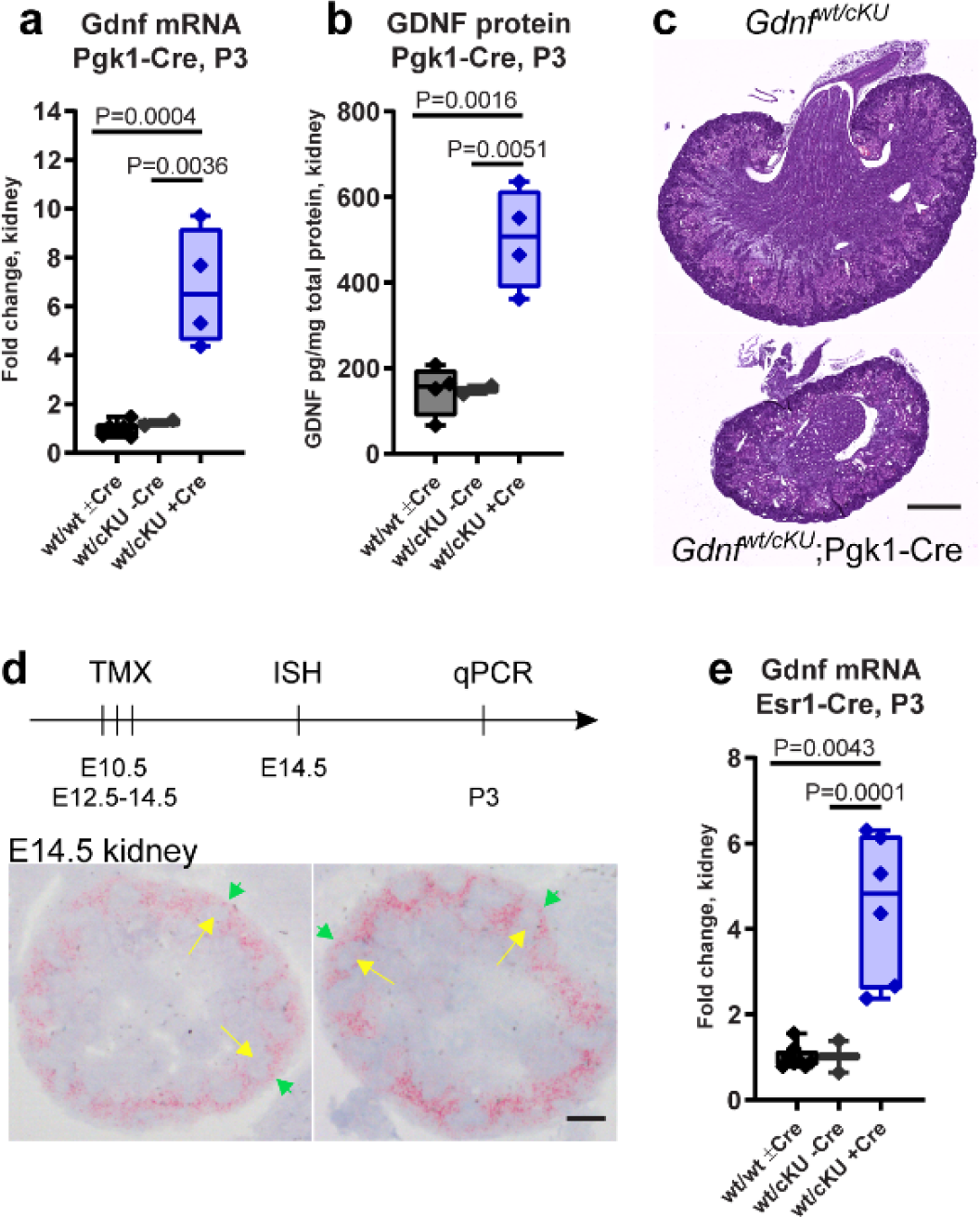
Endogenous GDNF in the kidney. **a-b** Gdnf mRNA (a) and protein (b) levels in P3 kidneys. **c** Kidney hematoxylin-eosin staining at P3. **d** Timeline of tamoxifen injection (top panel) and representative *in situ* hybridization image (bottom panel) from E14.5 kidney. Green arrows indicate GDNF-expressing metanephric mesenchyme and yellow arrows indicate ureteric buds. **e** Gdnf mRNA expression in P3 kidneys after tamoxifen injection. Median, upper and lower quartiles, and maximum and minimum values are shown. Scale bar 500 μm (c) or 100 μm (d). See Methods for details.

Next, we crossed *Gdnf*^*cKU*^ mice to a tamoxifen-inducible Esr1-Cre line^23^ and administered tamoxifen to pregnant females at E10.5. At E14.5, *in situ* hybridization using RNAscope probes against Gdnf revealed a stronger signal in *Gdnf*^*wt/cKU*^;Esr1-Cre mice compared to wild-type littermates, whereas Gdnf’s expression pattern in these mice was normal (Fig. 2d). To measure Gdnf mRNA levels, tamoxifen was then injected on three consecutive days at E12.5-E14.5, and mice were dissected at P3. We found that Gdnf expression in the kidneys of *Gdnf*^*WT/cKU*^;Esr1-Cre mice was increased by more than 4-fold compared to wild-type and *Gdnf*^*WT/cKU*^ controls (Fig. 2e). Therefore, we conclude that the *Gdnf*^*cKU*^ allele allows us to increase endogenous *Gdnf* expression at different stages of development, and, in line with our hypothesis, does not change normal *Gdnf* expression pattern.

### *Gdnf*^*cKU*^ mice have schizophrenia-like features

We next crossed the *Gdnf*^*cKU*^ mice to a Nestin-Cre-expressing mouse line^24^ to restrict the increase in endogenous GDNF levels to cells expressing GDNF in the CNS from E11.5 onwards. In the adult nigrostriatal system, GDNF is mostly produced in a sparse population of striatal interneurons^25^ that regulates striatal output^26^. We analysed *Gdnf* expression in the striatum of *Gdnf*^*cKU*^;Nestin-Cre mice and found that both Gdnf mRNA and GDNF protein levels were increased in *Gdnf*^*cKU*^ mice (Fig. 3a and Extended Data Fig. 2a), whereas Gdnf’s expression pattern appeared normal (Fig. 3b). In line with previous findings^7^, *Gdnf*^*cKU*^;Nestin-Cre mice exhibited increased levels of total tissue dopamine in the striatum and substantia nigra (SN) (Fig. 3c-d) and increased evoked dopamine release and uptake in the striatum (Fig. 3e-f).

**Figure 3.**
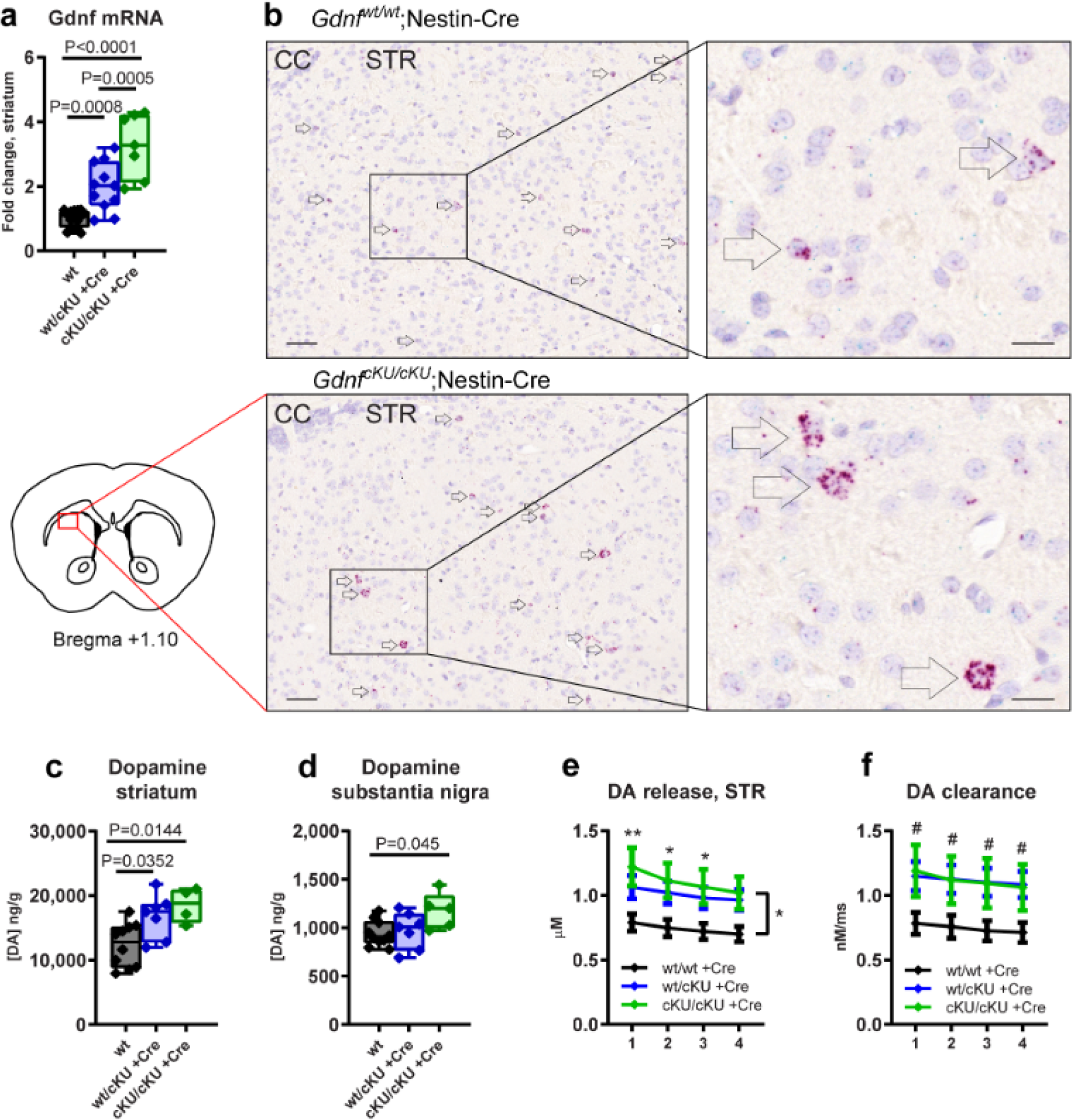
Endogenous GDNF in the brain of adult *Gdnf*^*cKU*^;Nestin-Cre mice. **a-b** Gdnf mRNA levels (a) and expression pattern revealed by RNAscope probes (b) in the striatum. **c-d** Total tissue dopamine levels in the striatum (c) and SN (d). **e-f** Stimulated dopamine release (e) and re-uptake (f) in striatal slices measured with CV. **a, c-d** Median, upper and lower quartiles, and maximum and minimum values are shown. **e-f** Error bars denote mean ± SE. Scale bar 50 μm (b, left panel) or 10 μm (b, right panel). See Methods for details.

Increased striatal dopamine signalling, particularly in the dorsal striatum, is a hallmark of schizophrenia^14–16^. Furthermore, studies on schizophrenic patients and a mouse model overexpressing dopamine receptor D2 (*Drd2*) have suggested that increased dopamine signalling in the striatum results in reduced dopamine signalling in the prefrontal cortex (PFC)^18,19^. We therefore asked if increased dopamine signalling in the striatum of *Gdnf*^*cKU*^;Nestin-Cre mice alters dopamine levels in the prefrontal cortex. Indeed, we found that dopamine levels were markedly reduced in the PFC of *Gdnf*^*wt/cKU*^;Nestin-Cre and *Gdnf*^*cKU/cKU*^;Nestin-Cre mice at two months of age (Fig. 4a), indicating that an increase in presynaptic dopamine in the nigrostriatal system also leads to aberrant dopamine signalling in the PFC in our mouse model.

**Figure 4.**
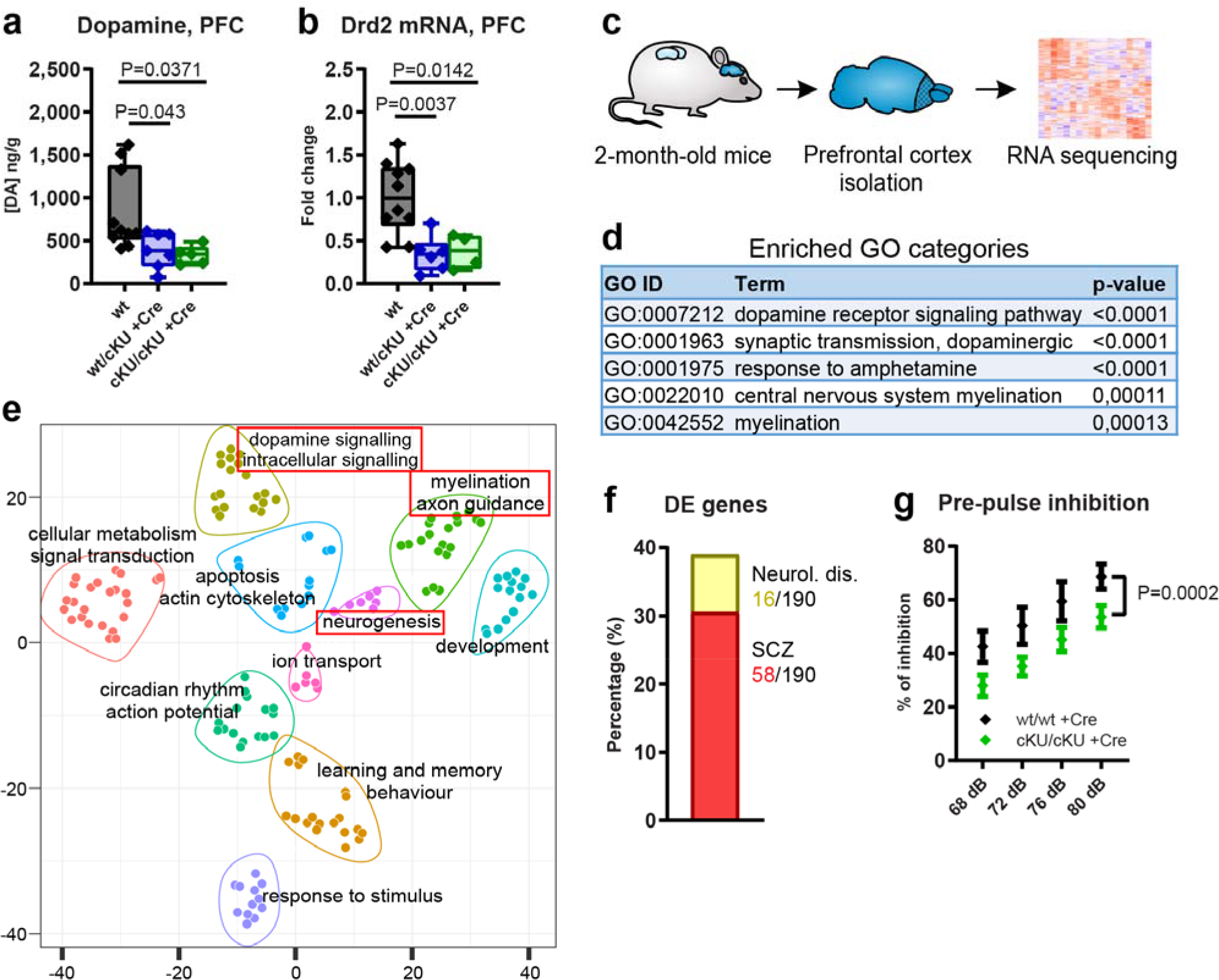
Schizophrenia-related phenotypes in *Gdnf*^*cKU*^;Nestin-Cre mice. **a** Dopamine levels in the PFC. **b** Drd2 mRNA levels. **c** Schematic of RNA sequencing experiment. **d** Enriched GO categories in *Gdnf*^*cKU/cKU*^;Nestin-Cre (homozygous) mice. **e** Scatter plot of GO enrichment analysis in *Gdnf*^*cKU/cKU*^;Nestin-Cre mice. Gene ontology categories most associated with schizophrenia are highlighted with red boxes. **f** Proportion of DE genes associated with schizophrenia or other neurodevelopmental disorders. **g** Pre-pulse inhibition test. GO, Gene Ontology. **a-b** Median, upper and lower quartiles, and maximum and minimum values are shown. **g** Error bars denote mean ± SE. See Methods for details.

Studies on patients with schizophrenia have identified several changes in gene expression in the PFC. In particular, reduced *Drd2* expression in the PFC is associated with schizophrenia^27–30^. We found that Drd2 mRNA levels were significantly reduced in the PFC of *Gdnf*^*wt/cKU*^;Nestin-Cre (heterozygous) mice and *Gdnf*^*cKU/cKU*^;Nestin-Cre (homozygous) mice (Fig. 4b). To obtain a more in-depth understanding of gene expression changes in the PFC, we performed RNA sequencing (Fig. 4c). We identified 311 differentially expressed (DE) genes in *Gdnf*^*wt/cKU*^;Nestin-Cre mice and 214 genes in *Gdnf*^*cKU/cKU*^;Nestin-Cre mice (Extended Data Fig. 2b-d). Functional annotation highlighted genes belonging to Gene Ontology (GO) categories related to dopamine signalling and myelination as most enriched among DE genes (Fig. 4d, Extended Data Fig. 2e). Further analysis on the top 150 enriched GO categories revealed 10 distinct clusters of DE genes, mostly related to nervous system development and function (Fig. 4e). Moreover, we found that many of the DE genes have previously been associated with schizophrenia in either genome-wide association studies or based on gene expression changes in schizophrenic patients (Fig. 4f and Extended Data Table 1), suggesting that the dysregulated pathways in *Gdnf*^*wt/cKU*^;Nestin-Cre and *Gdnf*^*cKU/cKU*^;Nestin-Cre mice share similarities with schizophrenia.

To investigate the behavioural outcome of aberrant dopamine signalling and gene expression in the PFC of *Gdnf*^*cKU/cKU*^;Nestin-Cre mice, we used the pre-pulse inhibition (PPI) test, which reflects the ability to successfully integrate and inhibit sensory information. Patients with schizophrenia and animal models with schizophrenia-like features have diminished responses to PPI^18,19,31–33^, which is believed to reflect impaired striatal and cortical dopamine transmission^32–34^. We found that the extent of PPI was significantly reduced in *Gdnf*^*cKU/cKU*^;Nestin-Cre mice (Fig. 4g), suggesting that altered dopamine signalling is paralleled by a deficit in sensorimotor gating. Therefore, we conclude that a tissue-specific increase in endogenous GDNF expression in the CNS at midgestation results in a phenotypic resemblance to schizophrenic patients, suggested by increased presynaptic dopamine signalling in the striatum, reduced dopamine levels in the PFC, reduced expression of several genes and pathways identified in post-mortem samples from the PFC of schizophrenic patients, and a deficit in PPI, a known endophenotype of schizophrenia (Table 1).

**Table 1.** Phenotypes of *Gdnf*^*cKU/cKU*^;Nestin-Cre mice that resemble findings from patients with schizophrenia.

| Phenotype | $Gdnf^{cKU/cKU};$<br>Nestin-Cre | $Gdnf^{cKU/cKU}$<br>AAV-Cre | Findings in<br>humans |
| --- | --- | --- | --- |
| *Increased striatal DA levels* | + | +/- | +<sup>15</sup> |
| *Reduced prefrontal cortex DA levels* | + | + | +<sup>27</sup> |
| *Reduced PFC dopamine D2 receptor levels* | + | n.a | +<sup>27-30</sup> |
| *Deficit in pre-pulse inhibition* | + | — | +<sup>35,36</sup> |

### Adult-onset GDNF cKU affects PFC DA system

Given that schizophrenia symptoms usually start in adolescence or adulthood, we were interested in whether the schizophrenia-like features found in *Gdnf*^*wt/cKU*^;Nestin-Cre and *Gdnf*^*cKU/cKU*^;Nestin-Cre mice could also be induced upon an adult-onset increase of endogenous GDNF expression. To that end, we injected AAV-Cre bilaterally into the striata of adult *Gdnf*^*wt/cKU*^ and *Gdnf*^*cKU/cKU*^ mice (Fig. 5a). We found that 2 months after AAV-Cre injection *Gdnf* expression was increased by more than 2-fold (Fig. 5b), resulting in an increase in dopamine release in response to a burst stimulus (Extended Data Fig. 3a). At the same time, total levels of striatal dopamine were not significantly increased, and evoked dopamine release and uptake were not affected (Extended Data Fig. 3b-d). However, dopamine levels in the PFC were significantly reduced by ∼50% (Fig. 5c), suggesting that changes in PFC dopamine signalling can be triggered already by 2-fold increase in endogenous GDNF levels in the adult striatum and by relatively minor changes in striatal dopamine function. We also administered the PPI test but found no indication of deficits in sensorimotor gating at the tested time point (Fig. 5d).

**Figure 5.**
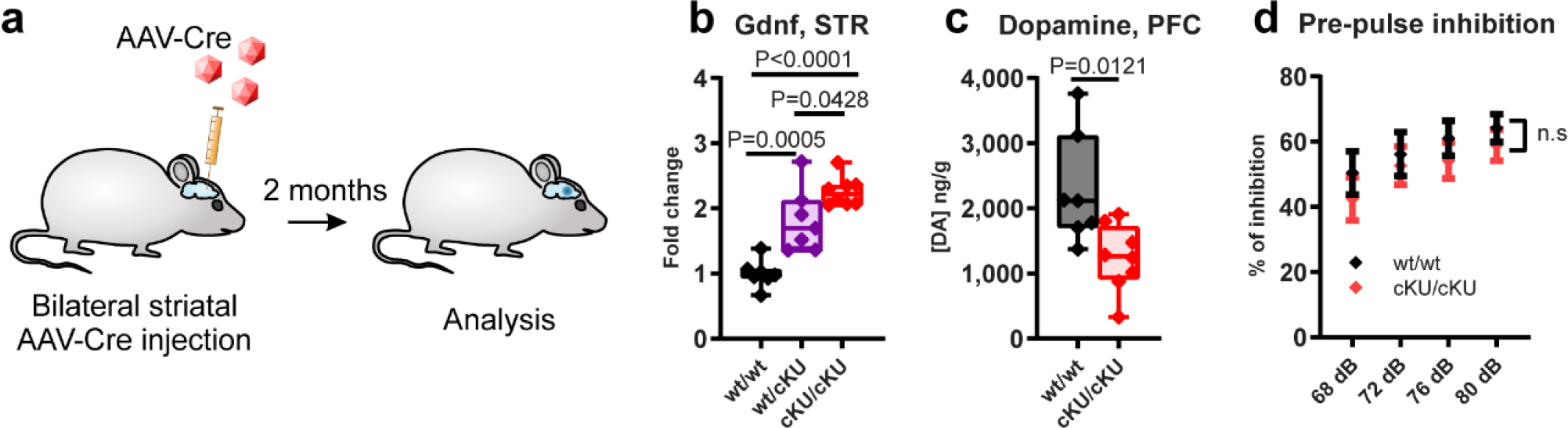
Dopamine system in *Gdnf*^*cKU*^ mice after bilateral striatal AAV-Cre injection. **a** Experiment schematic. **b** Striatal Gdnf levels 2 months after AAV-Cre. **c** Total levels of dopamine in the PFC. **d** Pre-pulse inhibition test. **b-c** Median, upper and lower quartiles, and maximum and minimum values are shown. **d** Error bars denote mean ± SE See Methods for details.

Taken together, we propose that *Gdnf*^*cKU*^ mice may present a useful tool for studying the pathologies and treatments arising from increased presynaptic striatal dopamine function, as well as for finding therapeutic strategies that specifically target this deficit in schizophrenia. The ability to upregulate endogenous gene expression in developing, adolescent and adult animals may also allow researchers to investigate the importance of timing the onset of altered protein levels, which could have relevance for certain nervous system diseases^37,38^.

### 3’UTR editing with CRISPR-Cas9

Conditional replacement of the 3’UTR is possible in experimental animals, but not applicable for tissue engineering or potential therapeutic uses. Based on the above findings on the conditional 3’UTR replacement, we hypothesised that removing inhibitory sequences from the endogenous 3’UTR via 3’UTR editing using the CRISPR-Cas9 system could result in the elevation of endogenous gene expression *in vivo* in wild-type animals.

To test this hypothesis, we decided to delete ∼2.5 kb of the Gdnf 3’UTR central region from the *Gdnf* gene, which contains negative regulation sequences that destabilize Gdnf mRNA^7^. Guide RNAs (gRNAs) were designed to target the beginning and end of the genomic region corresponding to the Gdnf 3’UTR, removing most of the predicted microRNA (miR) binding sites, but leaving the translational stop signal and polyadenylation signal intact [*Gdnf* <u>K</u>nock <u>U</u>p (KU) allele, Fig. 6a]. We hypothesised that the resulting transcript carrying a short 3’UTR with few miR binding sites would be more stable and lead to increased GDNF levels.

**Figure 6.**
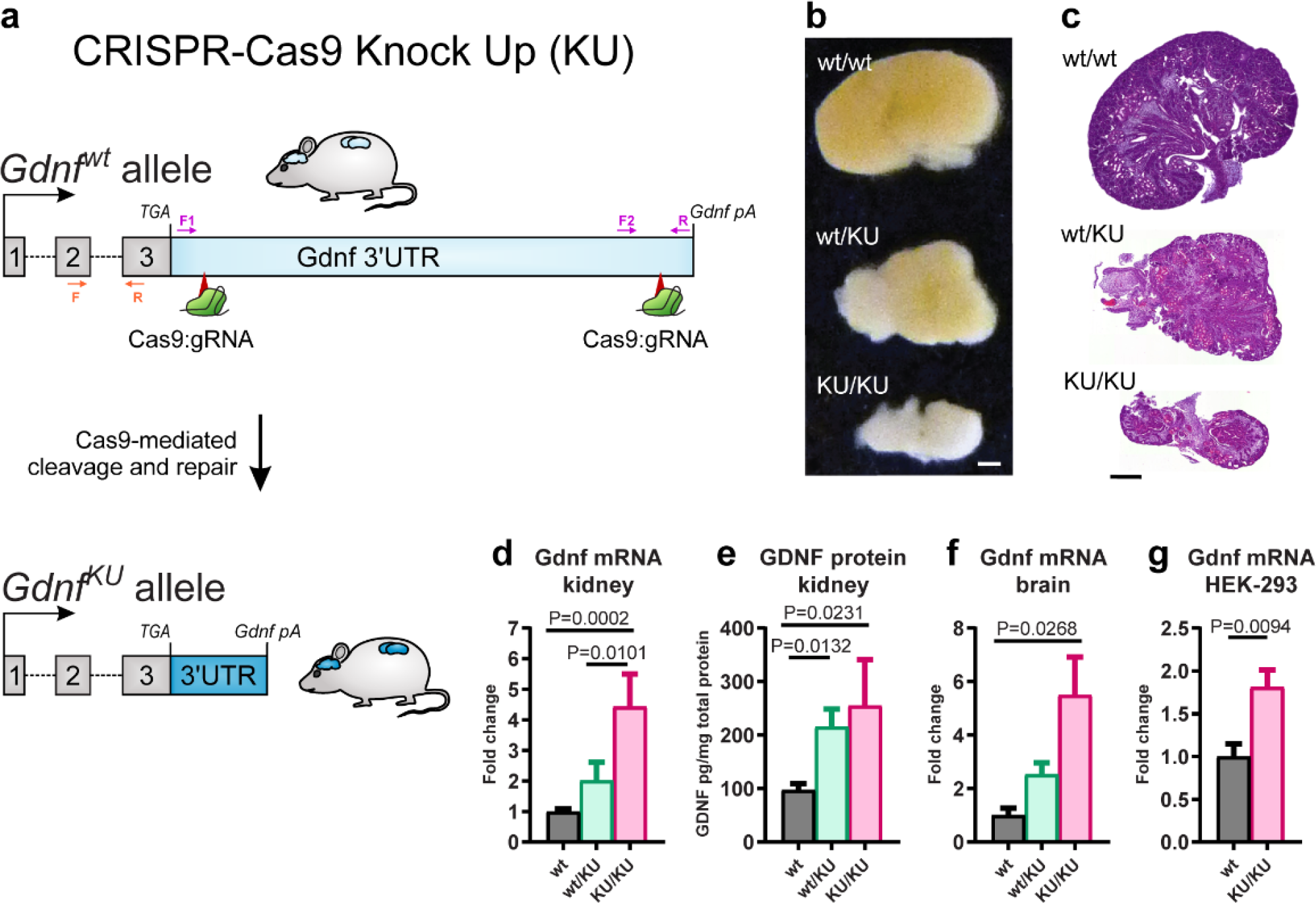
3’UTR editing with CRISPR-Cas9. **a** Schematic of gene <u>K</u>nock <u>U</u>p (KU) via 3’UTR editing. Locations of genotyping and qPCR primers are shown in purple and orange, respectively. **b-c** Representative images of gross morphology (b) and histology (c) of E18.5 kidneys. **d-e** Gdnf mRNA (d) and protein (e) levels in the kidney. **f** Gdnf mRNA levels in the brain. **g** Gdnf mRNA levels in HEK-293 cells after 3’UTR editing. Error bars denote mean ± SE. Scale bar 500 μm. See Methods for details.

Mouse zygotes were microinjected at the one-cell stage with purified gRNAs and *S. pyogenes* Cas9 mRNA, and embryos were dissected at E18.5 for analysis of the kidneys and brain. In embryos carrying one or two *Gdnf*^*KU*^ alleles (for detailed description on genotyping, please see the Methods section), the kidneys were notably smaller than kidneys from control animals (Fig. 6b) and, similar to kidneys from GDNF hypermorphs^7^ and *Gdnf*^*wt/cKU*^;Pgk1-Cre mice (Fig. 2c), displayed disorganized cortex-medulla structure (Fig. 6c). In line with these results, both Gdnf mRNA and GDNF protein levels were allele-dose-dependently increased in the kidneys of *Gdnf*^*wt/KU*^ and *Gdnf*^*KU/KU*^ mice (Fig. 6d-e). Furthermore, we found that Gdnf mRNA levels were also increased in the brains of *Gdnf*^*KU/KU*^ mice (Fig. 6f). Finally, we found that 3’UTR editing increased Gdnf mRNA expression in the human embryonic kidney (HEK-293) cell line (Fig 6g), suggesting that elevating endogenous gene expression via 3’UTR editing is also applicable to human cells.

Taken together, our data suggest that excising a long 3’UTR fragment from the *Gdnf* gene increases GDNF expression in human cells and mice. Importantly, the change in GDNF levels was achieved in human cells and wild-type mice without the introduction of a foreign sequence into the genome of the animals, suggesting that endogenous gene expression can also be elevated by editing the native 3’UTR of the gene. Therefore, this approach may be useful for identifying disease pathways in *in vitro* models and adult animals, as well as for developing treatment strategies for human disease.

## Discussion

Currently, only 1-2% of the entire human proteome can be targeted by pharmacological agonists or activators, with the majority of targets being receptors, enzymes and transporter proteins^39,40^. Many of the drugs have more than one target, making it difficult to pinpoint, which of the targets are responsible for treatment or for side-effects. Therefore, a tool that enables researchers to specifically Knock Up potential drug targets one at a time would assist in identifying endogenous pathways where upregulation is advantageous in disease models.

Elevation of endogenous protein expression due to copy number variations or mutations that alter regulatory sequences of genes contributes to many human diseases, including cancers, Alzheimer’s disease, Parkinson’s disease, autism and schizophrenia^37,41–45^. Using the neurotrophic factor GDNF as a proof of concept example, we show that endogenous gene Knock Up is possible both by conditionally replacing the 3’UTR with a sequence not targeted by inhibitory factors, and by cleaving the negative regulatory regions of the native 3’UTR using the CRISPR-Cas9 system. We find no changes in cell- and developmental stage specific expression pattern of *Gdnf*, consistent with the notion that the Knock Up approach does not alter the chromatin context of the gene. This feature makes gene Knock Up different from transgenic overexpression, where epigenetic regulation of transgene expression is often affected by the integration site of the transgene. Furthermore, gene Knock Up also differs from transcriptional activation where a nuclease dead endonuclease is fused with a transcriptional activation domain^46^, because the latter, similarly to transgenic overexpression, is likely to induce ectopic gene expression in cells where the gene is not normally expressed.

We found that CNS-restricted elevation of endogenous GDNF led to several phenotypes associated with schizophrenia, including increased presynaptic dopamine levels in the striatum and SN, reduced dopamine levels in the PFC, associated changes in gene expression in the PFC, and impaired sensorimotor gating. Furthermore, adult-onset increase in striatal GDNF expression also reduced dopamine levels in the PFC. Several genetic association studies have implicated a link between *GDNF* and schizophrenia, although the results are inconclusive as yet (Extended Data Table 2). Our data suggests that a change in endogenous GDNF levels at a specific stage in development is sufficient to drive schizophrenia-like phenotypes in mice, hinting at the possibility that aberrant regulation of *GDNF* expression may also lead to detrimental consequences in humans.

Importantly, while GDNF Knock Up in the brain resulted in altered dopamine levels in both the striatum and prefrontal cortex, conditional knockdown of GDNF expression with the same Nestin-Cre mouse line did not alter total tissue dopamine levels in the PFC^47^. Therefore, the results of this study emphasise that downregulation and upregulation of endogenous gene expression do not always lead to opposite phenotypes, and thus both approaches are necessary to understand gene function.

Our results also illustrate how an increase in endogenous gene expression levels at different developmental ages can lead to different phenotypic outcomes. Mice with a constitutive increase in endogenous GDNF levels (*Gdnf*^*wt/hyper*^)^7^ and mice with constitutively active GDNF receptor RET proto-oncogene signalling (*Ret*^*MEN2B/MEN2B*^)^48^ have increased striatal dopamine levels, but no change in PFC dopamine levels^49^ (Extended Data Fig. 3e) and no deficit in PPI^50^. In accordance with this finding, patients with multiple endocrine neoplasia, type 2B (MEN2B), expressing constitutively active RET have not been reported to display neurological deficits or schizophrenia. At the same time, PFC dopamine levels in adult *Gdnf*^*cKU/cKU*^ mice are dramatically reduced two months after endogenous GDNF upregulation in the striatum even with minor changes in striatal dopamine levels (Fig 5a-d). Therefore, the time of onset of an increase in endogenous gene expression clearly affects brain development and function, and methods like gene conditional Knock Up can assist in uncovering the function of endogenous genes at specific developmental time points.

The levels of more than one half of all protein-coding genes are predicted to be negatively regulated through the 3’UTR^51,52^. Therefore—as also supported by results from a recent *in vitro* study^53^—3’UTR editing could be used to upregulate the expression of many or most of these genes in their native chromatin context. At the same time, it is increasingly recognised that 3’UTRs can have other functions, such as regulating mRNA localization or protein-protein interactions^54–56^, which are likely to limit the use of the 3’UTR editing for gene Knock Up. However, once relevant motifs are identified, it could be feasible to retain the sequences required for these other functions by selecting appropriate gRNAs for Cas9-mediated excision, or by incorporating those motifs into the FLEx cassette, thus expanding the list of potential target genes for the Knock Up approach.

Taken together, we believe that the described Knock Up modes: constitutive (Pgk1-Cre), tissue specific (Nestin-Cre), inducible (Esr1-Cre), adult-onset (AAV-Cre) and CRISPR-Cas9-driven Knock Up in wild-type mice each provide new opportunities for both basic research and translational medicine.

## Methods

### Animals

All animal experiments were conducted according to the 3R principles of the European Union Directive 2010/63/EU governing the care and use of experimental animals, and following local laws and regulations (Finnish Act on the Protection of Animals Used for Scientific or Educational Purposes (497/2013), Government Decree on the Protection of Animals Used for Scientific or Educational Purposes (564/2013)). The protocols were authorised by the national Animal Experiment Board of Finland (license numbers ESAVI/11198/04.10.07/2014 and ESAVI/12046/04.10.07/2017). Mice were maintained in a 129Ola/ICR/C57bl6 mixed genetic background. The mice were group-housed in a specific pathogen free stage with *ad libitum* access to food and water under a 12-h light-dark cycle (lights on at 6 a.m.) at relative humidity of 50-60 % and room temperature 21±1 °C. Bedding (aspen chips, Tapvei) and nest material (Tapvei) were changed weekly and wooden blocks (Tapvei) were provided for enrichment. All behavioural tests, tissue collection and processing were performed by researchers blinded to the genotypes of the animals and in all experiments, wild-type littermates were used as controls.

### Generation of *Gdnf*^*cKU*^ knock-in allele

The targeting construct for the *Gdnf*^*cKU*^ allele (S Fig 1A) contained a 4021 bp 5’ homologous arm spanning part of the second intron and the entire coding sequence of the mouse *Gdnf* gene, a 610 bp cassette containing the bovine growth hormone polyadenylation signal (bGHpA) in an inverted orientation flanked by the FLEx system^17^ starting immediately after the stop codon, a 2615 bp Pgk1-puΔtk-bGHpA sequence flanked by Frt sites, and a 2927 bp 3’ homologous arm. The targeting vector was synthesized by GenScript (NJ, USA) and confirmed by restriction analysis and sequencing. G4 embryonic stem cells (derived from mouse 129S6/C57BL/6Ncr) were cultured on puromycin-resistant primary embryonic fibroblast feeder layers, and ∼10^6^ cells were electroporated with 30 μg of linearized targeting construct. After electroporation the cells were plated on 10-cm culture dishes and exposed to puromycin (1.5 μg/ml; Sigma). Colonies were picked after 7-9 days of selection and grown on a 96-well plate. DNA isolated from ES cells was screened by long-range PCR for both 5’ and 3’ homologous arms. In total, 480 clones were screened. Correct PCR products were verified by sequencing. Correctly targeted ES cells were injected into C57BL/N6 (Charles River Laboratories, MA, USA) mouse blastocysts to generate chimeric mice. Germline transmission was achieved by breeding male chimeras with C57BL/N6 females. The Pgk1-puΔtk-bGHpA sequence was removed using CAG-Flp mouse line at F2 generation.

### AAV production

Pseudotyped AAV2/5 vectors were produced using a double-transfection method with the appropriate transfer plasmid and the helper plasmid containing the essential adenoviral packaging genes, as described previously^57^. Vectors were purified by iodixanol step gradients and Sepharose Q column chromatography. The purified viral vector suspension was titrated with TaqMan quantitative PCR and primers targeting the WPRE sequence.

### Stereotaxic surgery

Mice were anesthetized with isoflurane (3-4% for induction and 1.5-2% for maintenance; Oriola, Finland) in 100% oxygen. The hair on the top of the head was shaved and cleaned with Desinfektol P (Berner Pro, Finland). Mice were placed on a stereotaxic apparatus and lidocaine (Yliopiston Apteekki, Finland) was injected into the skin for local analgesia. The skin was cut with a scalpel and holes were drilled into the skull bilaterally. Two different sets of coordinates were used throughout the study: 1) A/P 1.2, M/L ±2.2 mm and A/P 0.2, M/L ±2.2 mm, or 2) A/P 0.7, M/L ±2.2, relative to the Bregma. With respect to virus diffusion and *Gdnf* expression, both injection regimens produced a comparable outcome. A 33G blunt NanoFil needle (World Precision Instruments, USA) was inserted into the brain parenchyma at a 10-degree angle until reaching D/V −3.0 mm, relative to Bregma. 1.0 μl (injections according to the first set of coordinates) or 1.5 μl (injections according to the second set of coordinates) of AAV5-Cre in Dulbecco’s PBS, corresponding to 1.7 × 10^11^ viral genome copies, was injected into each site at a flow rate of 0.2 μl/min. The needle was kept in place for an additional 4-5 minutes and retracted slowly. The skin was closed with sutures and animals were subcutaneously injected with 5 mg/kg Rimadyl or Norocarp (Yliopiston Apteekki, Finland) in saline for post-operative analgesia. The mice were 3-5 months old at the time of the injections. Wild-type littermates were used as controls and were injected with the same dose of AAV5-Cre as *Gdnf*^*cKU*^ mice.

### Tamoxifen injection

For the analysis of Gdnf mRNA and protein levels, *Gdnf*^*wt/cKU*^ mice were crossed to Esr1-Cre mice and pregnant females were injected i.p. with 25 mg/kg tamoxifen (Sigma-Aldrich) in corn oil on three consecutive days between E12.5-E14.5. The pups were dissected at P3 and tissues were isolated for further analysis. For RNAscope analysis, pregnant females were injected with 25 mg/kg tamoxifen at E10.5 and tissues were harvested at E14.5.

### Pre-pulse inhibition

The PPI test was performed as described previously^50^. Male *Gdnf*^*cKU*^;Nestin-Cre mice were analysed at an age of 3-4 months by experimenters blinded to the genotypes of the mice. Littermates were used as controls.

### gRNA design and production for zygote microinjection

Guide RNAs were designed using the CRISPR Design Tool (crispr.mit.edu)^58^ and produced using overlap PCR^59^. The gRNA template was transcribed *in vitro* with the Megashortscript T7 Kit (AM1354, Life Technologies) by incubating 165 ng of template for 5 hours at 37 °C. *In vitro* transcribed RNA was purified with Megaclear Kit (AM1908, Life Technologies) and gRNA integrity was measured with Agilent 2100 Bioanalyzer (Agilent Technologies).

Two gRNAs were designed to target both the beginning (A, B) and the end (C, D) of the 3’UTR, while leaving the translational stop signal and putative polyadenylation signal intact (Fig. 6a). All of the possible combinations (A+C, A+D, B+C and B+D) were used for zygote injections and the efficiencies of different combinations were comparable to each other (not shown). gRNA target sequences are shown in Extended Data Table 3.

### Zygote microinjection

Cas9 mRNA (5meC, Ψ, L-6125, Trilink Biotechnologies) and purified gRNAs were injected into the pronucleus at concentrations of 25 ng/ml and 12.5 ng/ml, respectively, in an injection buffer containing 0.25 mM EDTA and 10 mM Tris-HCl, pH 7.4. Prior to microinjection, the mixture was centrifuged at 13,200 rpm for 30 min at 4 °C in a standard table top centrifuge and the supernatant was collected into a fresh RNase-free tube.

### Isolation of tissues

Pregnant females were deeply anesthetized with CO_2_, followed by cervical dislocation and decapitation, and embryos were dissected at indicated time points. P3 mice were sacrificed by decapitation. The kidneys were either immersed in 4% formaldehyde (Sigma, in 1× PBS) overnight, followed by dehydration and paraffinization (Leica ASP 200), or immediately snap frozen on dry ice and stored at −80 °C until analysed for mRNA and protein expression. Brains from E18.5 embryos were snap frozen and stored at −80 °C.

To isolate samples for biochemical analysis, adult mice were deeply anesthetized with CO_2_, followed by cervical dislocation and decapitation. The brain was quickly dissected and immersed in ice-cold PBS. Brain regions of interest were isolated from 2-mm slices cut using a brain matrix (Stoelting). Tissue samples dedicated for RNA extraction or HPLC, as well as for kidney GDNF ELISA, were immediately snap frozen on dry ice and stored at −80 °C until processed. For GDNF ELISA, brain samples were immediately lysed in lysis buffer (see below) and stored at −80 °C until analysis.

To isolate samples for *in situ* hybridization, adult mice were anesthetized with pentobarbital (Mebunat, 200 mg/kg, i.p., Yliopiston Apteekki) and perfused with warm PBS and 4% formaldehyde. Samples were post-fixed in 4% formaldehyde for 24 hours at room temperature and paraffinised or dehydrated in 30% sucrose (Thermo Fisher Scientific) in PBS prior to sectioning.

### RNA and protein isolation

Total RNA was isolated using Trizol Reagent (Thermo Fisher Scientific) or RNAqueous Micro Kit (Thermo Fisher Scientific). Tissue samples for protein analysis were homogenized in lysis buffer prepared according to the recipe in the GDNF Emax Immunoassay System protocol (Promega). The tissue homogenate was centrifuged at 5000 rpm for 15 minutes at 4 °C and the supernatant was used immediately or stored at −80 °C until further processing. Each protein sample was only thawed and used once.

### Genotyping

Genotyping samples were collected at weaning and/or during dissection. Genomic DNA was isolated from the tail using Extracta DNA Prep for PCR - tissue (Quanta Biosciences, USA). Routine genotyping was performed using AccuStart II GelTrack PCR SuperMix (Quanta Biosciences, USA) and analysed by electrophoresis using 1.5-2% agarose gels in borate buffer. Primers used for genotyping the mice for the presence of *Gdnf*^*cKU*^ allele are indicated on Extended Data Fig. 1a and Extended Data Table 3.

Genotyping of tissues obtained from 3’UTR editing using the CRISPR-Cas9 posed a challenge due to mosaicism in analysed tissues. Because GDNF is not expressed ubiquitously, but rather in defined cell types at various stages of development, we were unable to rely on genotyping from the genomic DNA. Instead, we used cDNA synthesised using random hexamer primers as a template for genotyping, to be able to specifically detect the presence of the cleaved allele in GDNF-expressing cells. To our surprise, the genotypes in the brain and kidney obtained from the same animals did not always coincide, suggesting that Cas9-mediated DNA cleavage and repair had taken place after the cellular lineages for the particular cell types had separated. Primers used for genotyping the *Gdnf*^*KU*^ allele are indicated in Extended Data Table 3 and Fig. 6a.

### Reverse transcription and quantitative PCR

150 to 500 ng of total RNA per sample were treated with RNase-free DNase I (Thermo Fisher Scientific). DNase I was inactivated with 5 mM EDTA at 65 °C, immediately followed by a reverse transcription reaction using random hexamer primers and RevertAid Reverse Transcriptase (Thermo Fisher Scientific). Complementary DNA (cDNA) was diluted 1:10 and stored at −20 °C until used for qPCR.

Quantitative PCR was performed with LightCycler 480 SYBR Green I Master (Roche) or with BioRad C1000 Touch Thermal Cycler upgraded to CFX384 System (BioRad), supplied with SYBR Green I Master (Roche) and 250 pmol primers, in 10 μl total volume in 384-well plates. Each reaction included cDNA or a negative control (minus-reverse transcription control or water). Mouse Actb or a combination of Pgk1, Hprt and Gapdh were used as reference genes. Primer sequences are given in Extended Data Table 3. All samples were analysed in duplicates. Results for a biological repeat were discarded when the C_q_ value for one or more of the replicates was 40 or 0, or when the C_q_ difference between replicates was >1.

### RNA sequencing

Total RNA was quality checked on an Agilent 2200 TapeStation system. RNA quantity was measured using a NanoDrop ND-1000 Spectrophotometer and 200 ng RNA/sample were used for library preparation using TruSeq Stranded mRNA protocol (Illumina). The yield and quality of the amplified libraries was analysed using Qubit (Thermo Fisher) and the Agilent 2200 TapeStation system. The indexed cDNA libraries were normalised and combined, and the pools were sequenced on the Illumina Nextseq for a 75-cycle v2.5 sequencing run, generating 75 bp single-end reads. Base calling and demultiplexing was performed using CASAVA software with default settings generating Fastq files for further downstream mapping and analysis.

Fastq files were aligned to the mouse reference genome (mm10) employing the STAR aligner^60^. Mapped reads from the subsequent bam files were counted in annotated exons using featureCounts^61^. Reference genome sequence in fasta format and entrez gene annotations were retrieved from UCSC. The resulting count table was imported to R/Bioconductor and differential gene expression analysis was carried out with edgeR^62^. Analysis was performed on genes that had 1 count per million (cpm) in at least 3 samples and normalised with TMM normalisation. Genes with an adjusted p-value under 0.05 were termed significant and were used for GO analysis using the topGO library^63^. A similarity matrix was constructed of the top 150 GO terms with the GOSemSim library^64^, which was then clustered by t-SNE. The t-SNE result was cut into clusters using dynamic tree cutting with the dynamicTreeCut library. Plots were prepared in ggplot2.

### Analysis of protein levels

Quantification of GDNF protein levels in the brain and kidney was performed using GDNF Emax Immunoassay (Promega) with acid treatment. Due to high background in protein samples derived from the striatum, a negative control sample from *Gdnf*^*cKO/KO*^;Nestin-Cre mice—dissected at the same time as experimental samples and obtained from an animal of comparable age—was always included in GDNF ELISA plate. We were not able to detect a difference between wild-type and *Gdnf*^*cKO/KO*^;Nestin-Cre samples if the brain tissue was snap frozen before lysis, therefore fresh tissue was always lysed before freezing and each frozen sample was thawed and used once. 20-100 μg of total protein from the brain or kidney, measured with the DC Protein Assay (Bio-Rad), was loaded on the ELISA plate. Signal obtained from the *Gdnf*^*cKO/KO*^;Nestin-Cre negative control sample was subtracted from experimental striatal samples. All samples were analysed in duplicates.

### HPLC

The levels of dopamine and its metabolites were analysed as previously described^65^, using HPLC with electrochemical detection.

### Histological analysis and *in situ* hybridization

For histological analysis, 5-μm paraffin sections were stained with Harris’s hematoxylin (Merck) and eosin Y (Sigma-Aldrich), according to a routine protocol. RNA *in situ* hybridization was performed as previously described^7^. Briefly, fresh 5-μm paraffin sections from embryonic kidney or adult brain were hybridized with RNAscope^66^ probes (Advanced Cell Diagnostics) detecting Gdnf mRNA, according to manufacturer’s recommendations. Sections were scanned with a Pannoramic 250 digital slide scanner (3D Histech).

### Cell culture

gRNAs targeting the genomic region corresponding to human Gdnf 3’UTR were designed using the CRISPR Design Tool (crispr.mit.edu)^58^ and are indicated in Extended Data Table 3. The gRNAs were cloned into pAAV-EF1a-eGFP^67^ (a gift from Brandon Harvey). Human embryonic kidney 293 (HEK-293) cells were cultured at 37 °C with 5 % CO_2_ in Dulbecco’s modified Eagle’s medium (DMEM) supplemented with 10 % fetal bovine serum (FBS; SV30160, Thermo Fisher Scientific, Waltham, MA, USA) and 1× Normocin (ant-nr-2, InvivoGen, San Diego, CA, USA). HEK-293 cells were passaged for two consecutive days before transfection with pAAV-dgRNA-EF1a-eGFP and SpCas9-2A-Puro (a gift from Feng Zhang, Addgene plasmid # 62988)^68^ plasmids using the calcium phosphate co-precipitation method. Medium was replaced 6 h after transfection, and cells were cultured for 48 h before sorting using BD Fluorescent Activated Cells Sorting (FACS) Aria II (BD Biosciences, San Jose, USA). After FACS, eGFP-positive cell population was plated in a 10-cm gelatinized plate (10^4^ cells/plate). Puromycin selection (1μg/ml) was maintained for 3 days, and single cell colonies were picked and expanded for DNA and RNA isolation.

### Fast-scan cyclic voltammetry

At 6-8 months of age, mice were decapitated, the brain was removed, and 300-μm-thick coronal slices containing the striatum were cut on a model 7000 smz-2 vibratome (Campden Instruments) in oxygen-bubbled (95% O2, 5% CO2) ice-cold cutting artificial cerebrospinal fluid (ACSF) containing the following (in mM): 75 NaCl, 2.5 KCl, 26 NaHCO3, 1.25 NaH2PO4, 2 MgCl2, 0.7 CaCl2, and 100 glucose. The slices were allowed to recover in a holding chamber for 20 min at 34°C, followed by 1-3 h at room temperature in oxygen-bubbled (95% O2, 5% CO2) recording ACSF containing the following (in mM): 119 NaCl, 3 KCl, 26 MgSO4, 1 KH2PO4, 1.2 MgCl2, 2 CaCl2, and 10 glucose. In the recording chamber, the slices were perfused continuously with oxygen-bubbled recording ACSF. Fast-scan cyclic voltammetry recordings were performed using cylindrical 5 μm carbon fiber electrodes positioned at the dSTR ∼50 μm below the exposed surface. Striatal slices were stimulated electrically at 2 min intervals using a stainless steel bipolar electrode placed ∼100 μm from the recording electrode. Square pulses of 0.4 ms (AAV-Cre injected mice) or 1 ms (*Gdnf*^*cKU*^;Nestin-Cre mice) in duration were produced using an Iso-Flex stimulus isolator triggered by a Master-8 pulse generator (A.M.P.I.). A stimulus magnitude of 200 mA was obtained by selecting the minimum value that produced the maximum response reliably. Triangular voltage ramps from a holding potential of −450 mV to +800 mV over 9 ms (scan rate of 300 mV/ms) were applied to the carbon fiber electrode at 100 ms intervals. In burst stimulus experiments, a burst of 5 pulses at 20 Hz was used. The current was recorded using an Axopatch 200B amplifier (Molecular Devices), filtered through a 5 kHz low-pass Bessel filter, and digitized at 40 kHz (ITC-18 board; InstruTech). Triangular wave generation and data acquisition were controlled and the recorded transients were characterized using a computer routine in IGOR Pro (WaveMetrics)^69,70^. Background-subtracted cyclic voltammograms obtained with 1 μM dopamine solution (dopamine-HCl; Sigma-Aldrich) were used to calibrate the electrodes.

### Statistical analyses

All values are presented as mean ± standard error of the mean. Statistical comparison between two groups was performed using unpaired Student’s *t*-test with two-tailed distribution. F-test was used to compare variances. Multiple comparisons were performed with one-way or two-way analysis of variance (ANOVA), followed by Tukey’s or Sidak’s *post hoc* test. Nonparametric data was analysed with Kruskal-Wallis test, followed by Dunn’s multiple comparison analysis. Statistical analysis was performed with GraphPad Prism v8. Quantitative PCR data was analysed as described previously^71^, using the geometric mean of reference genes for normalisation. Statistical significance level was set at *p* < 0.05. Details on statistical analysis are provided in Extended Data Table 4.

## Data availability

The GEO database accession number for the RNA sequencing data reported in this paper will be added here prior to publishing. Other datasets in the present study are available from the corresponding author on reasonable request.

## Acknowledgements

The authors wish to thank S. Wiss and J. Lahtinen for technical assistance and A. Damdimopoulus for bioinformatics analysis of RNA sequencing data. Genetically modified mice were generated at Turku Center for Disease Modeling, and the authors thank K. Hovirinta, N. Messner, M. Niiranen, H. Niittymäki and J. Palmu for technical assistance. K.M. was supported by the doctoral program Brain and Mind, Alfred Kordelin Foundation and Sigrid Juselius Foundation. J.O.A. was supported by the Academy of Finland (grant no. 297727), Sigrid Juselius Foundation, Alzheimerfonden, Faculty of Medicine at the University of Helsinki, Helsinki Institute of Life Science and by European Research Council (ERC) under the European Union’s Horizon 2020 research and innovation programme (grant no. 724922). G.T. was supported by the Finnish Parkinson Foundation. D.R.G. was supported by the Finnish Parkinson Foundation, Fulbright Finland and the Biomedicum Helsinki Foundation. A.P. was supported by Jane and Aatos Erkko Foundation. J.K. was supported by the Finnish Cultural Foundation and Emil Aaltonen Foundation. J.J was supported by the Swedish Research Council, the Swedish Foundation for Strategic Research, the Swedish Brain Foundation and the Swedish Government Initiative for Strategic Research Areas (MultiPark & StemTherapy).

## Author contributions

K.M. planned and designed experiments, analysed and interpreted the data, prepared figures and wrote the manuscript. K.M., S.O., D.R.G., A.M.R., G.T., L.L.P., A.P., T.P.P., J.K. and M.C.C. performed experiments. D.R.G. and A.P. designed and performed fast-scan cyclic voltammetry experiments. K.M. and N.S. analysed processed RNA sequencing data. F.Z. and P.S. generated the *Gdnf*^*cKU*^ allele. J.J. produced AAV. J.O.A. planned and designed experiments, interpreted the data, provided funding and contributed to writing the manuscript. All authors read and approved the manuscript.

