## Extended Data Figures for "Gene Knock Up via 3’UTR editing to study gene function *in vivo*"

Extended Data Figure 1, related to Figure 1

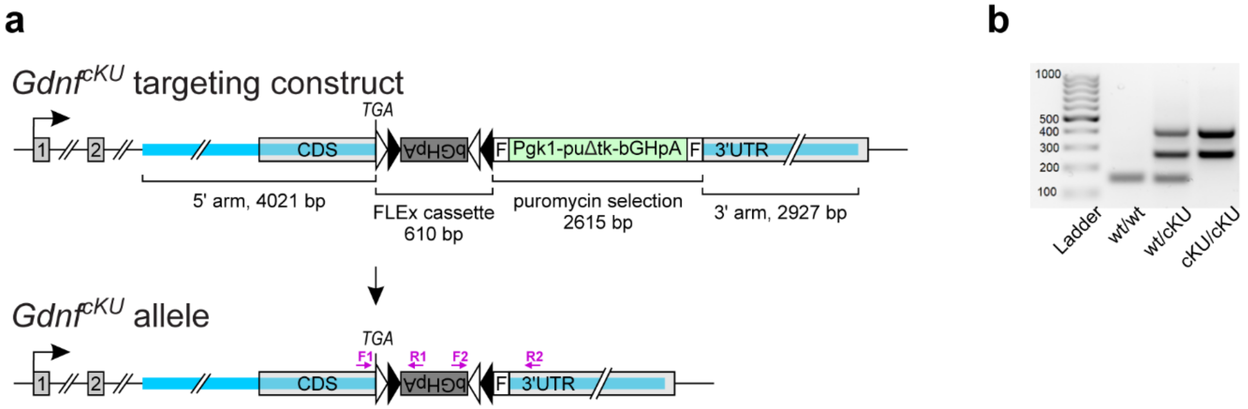


**Extended Data Figure 1** Generation of *Gdnf* conditional Knock Up (cKU) allele. **a** Targeting scheme (upper panel) showing 5' and 3' homologous arms (blue lines), FLEx cassette containing bovine growth hormone 3'UTR and polyadenylation signal (bGHpA) in an inverted orientation (grey box), and puromycin selection marker flanked with Frt sites (green box). Crossing to a Deleter Flp mouse line removes the selection cassette and results in the *Gdnf*$^{cKU}$ allele (lower panel). Purple arrows indicate the location of genotyping primers. **b** Representative genotyping image. CDS, coding sequence; F, Frt.

Extended Data Figure 2, related to Figures 3-4

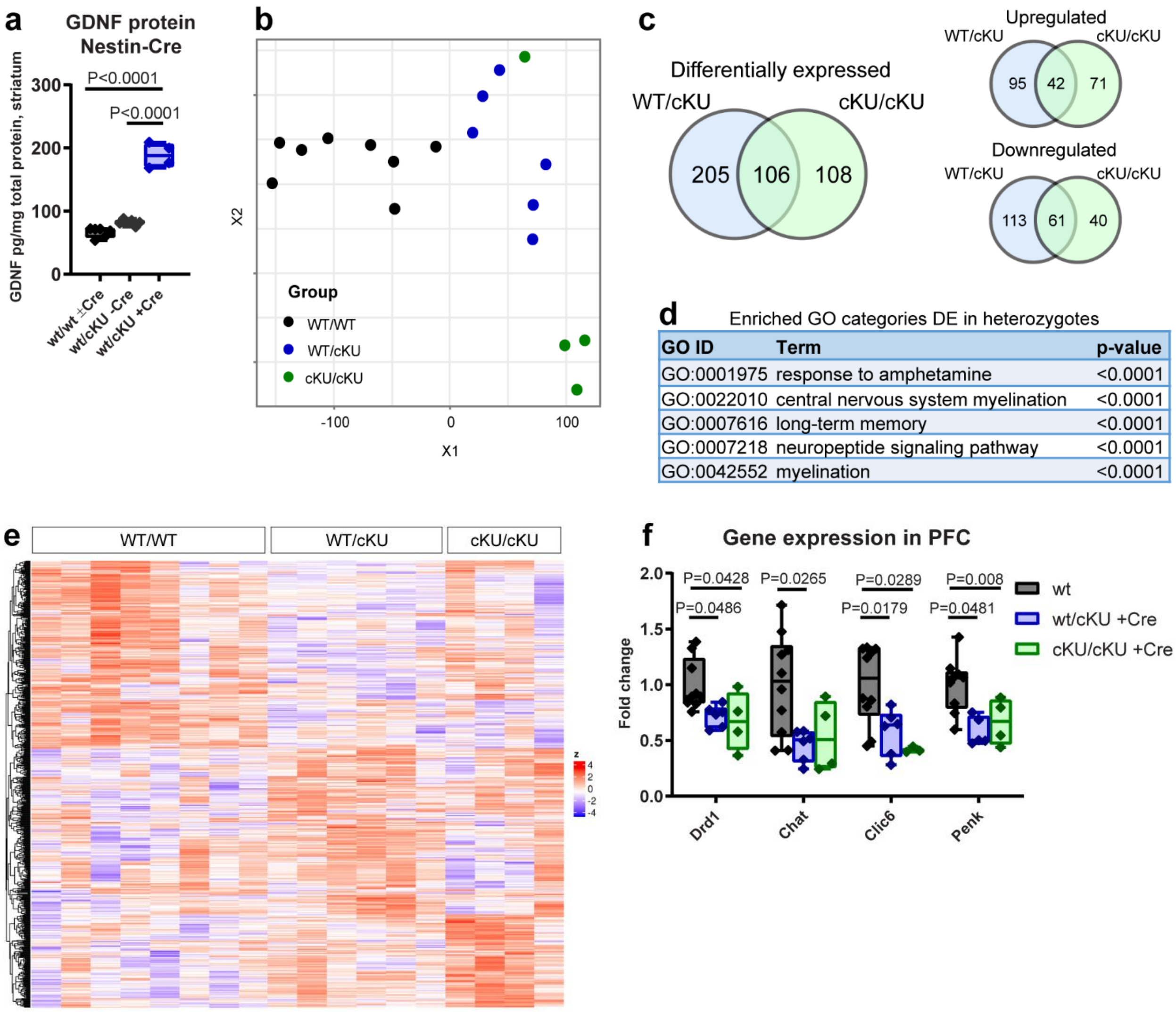


**Extended Data Figure 2** Dopamine system in *Gdnf*$^{cKU}$;Nestin-Cre mice. **a** GDNF protein levels in the striatum of adult *Gdnf*$^{cKU}$;Nestin-Cre mice. **b-e** RNA sequencing in the PFC of *Gdnf*$^{cKU}$;Nestin-Cre mice. **b** tSNE plot of RNAseq samples. **c** Heatmap of differentially expressed genes. **d** Venn diagram showing overlap of differentially expressed genes in *Gdnf*$^{wt/cKU}$;Nestin-Cre and *Gdnf*$^{cKU/cKU}$;Nestin-Cre mice**. e** GO categories enriched differentially expressed genes in *Gdnf*$^{wt/cKU}$;Nestin-Cre (heterozygous) mice. **f** Validation of the expression of selected differentially expressed genes using qPCR. Median, upper and lower quartiles, and maximum and minimum values are shown. See Methods for details.

Extended Data Figure 3, related to Figure 5

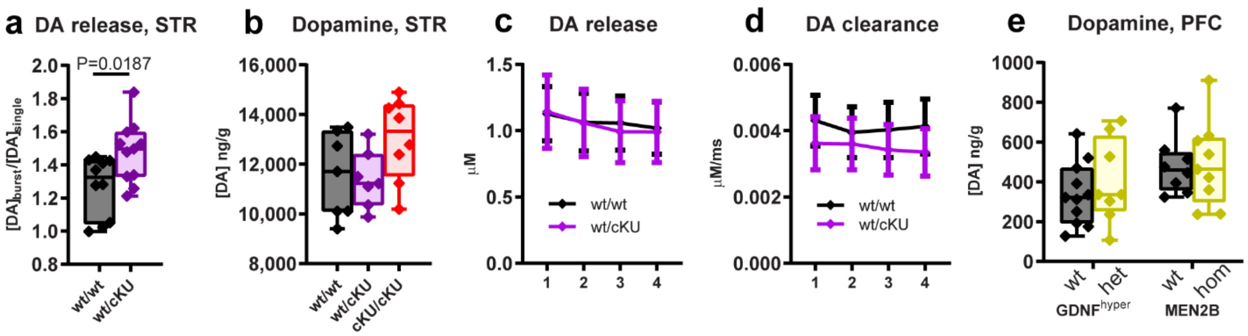


**Extended Data Figure 3** Dopamine system in *Gdnf*$^{cKU}$ mice two months after bilateral striatal AAV-Cre injection **a** Dopamine release in response to burst stimulus. **b** Total dopamine levels in the striatum. **c-d** Dopamine release (c) and re-uptake (d) in striatal slices from adult *Gdnf*$^{wt/wt}$ and *Gdnf*$^{wt/cKU}$ mice two months after striatal AAV-Cre injection. **e** Dopamine levels in the PFC of wild-type, *Gdnf*$^{wt/hyper}$ and *Ret*$^{MEN2B/MEN2B}$ (MEN2B) homozygous mice. **a-b, e** Median, upper and lower quartiles, and maximum and minimum values are shown. **c-d** Error bars denote mean ± SE. See Methods for details.
