## Extended Data Table 1 for "Gene Knock Up via 3’UTR editing to study gene function *in vivo*"

Mätlik *et al.* Gene Knock Up via 3'UTR editing to study gene function *in vivo*

Differentially expressed protein-coding genes in *Gdnf*$^{cKU/cKU}$;Nestin-Cre mice that have been associated with schizophrenia in mouse and/or human studies.

| *Gene symbol* | *P-value* | *Differential expression* | *Genetic association* | *Animal model* | *Reference* |
|---|---|---|---|---|---|
| *Adora2a* | <0.0001 | + | + | | 71-73 |
| *Adcy7* | 0,00032 | | + | | 74 |
| *Alas2* | 0,00048 | | + | | 75 |
| *Aldh1a1* | <0.0001 | + | + | | 73,76 |
| *Als2cl* | <0.0001 | | + | | 77 |
| *Ano2* | 0,00066 | | + | | 78 |
| *Arc* | <0.0001 | + | + | + | 73,79,80 |
| *Arnt* | 0,00015 | | + | | 81,82 |
| *Btg2* | <0.0001 | + | + | | 73,83 |
| *Cbln4* | 0,00026 | + | | | 73 |
| *Celsr3* | 0,00021 | | + | | 74 |
| *Cnp* | <0.0001 | + | | | 84 |
| *Col16a1* | <0.0001 | | + | | 74 |
| *Col19a1* | 0,00014 | | + | | 85 |
| *Cpeb1* | 0,00056 | | + | | 82,86 |
| *Cyr61* | <0.0001 | + | | | 73 |
| *Dennd6b* | <0.0001 | | + | | 74 |
| *Dnah6* | <0.0001 | | + | | 74 |
| *Drd1* | <0.0001 | + | + | | 81,87-91 |
| *Drd2* | <0.0001 | + | + | + | 72,81,87,92-99 |
| *Dusp1* | <0.0001 | + | | | 100 |
| *Egr1* | 0,00029 | + | + | | 73,92 |
| *Egr2* | <0.0001 | + | | | 73 |
| *Fos* | <0.0001 | | + | | 101 |
| *Galnt9* | <0.0001 | + | | | 73 |
| *Gpr88* | <0.0001 | | + | + | 102,103 |
| *Hr* | 0,00029 | + | | | 73 |
| *Ier2* | 0,00055 | | + | | 74 |
| *Il18bp* | 0,00053 | | + | | 104 |
| *Il3ra* | <0.0001 | | + | | 105-107 |
| *Inf2* | 0,00011 | + | | | 73 |
| *Lin28b* | <0.0001 | | + | | 86 |
| *Mag* | <0.0001 | + | + | | 108-111 |

| | | | | | |
|---|---|---|---|---|---|
| *Masp2* | 0,00074 | | + | | 74 |
| *Mbp* | <0.0001 | + | + | | 76,112,113 |
| *Mobp* | <0.0001 | + | + | | 76,111,114,115 |
| *Mog* | 0,00047 | + | + | | 111,116-119 |
| *Mov10* | 0,00015 | | + | | 74 |
| *Ndrg1* | 0,00045 | + | | | 73 |
| *Ndst3* | 0,00015 | | + | | 120-122 |
| *Npas4* | <0.0001 | + | | | 123 |
| *Nr4a1* | <0.0001 | + | | | 100,124 |
| *Pde10a* | <0.0001 | + | + | | 125,126 |
| *Pde7b* | <0.0001 | + | + | | 73,127,128 |
| *Peg3* | 0,00024 | | + | | 74 |
| *Penk* | <0.0001 | | + | | 129,130 |
| *Pkd1* | 0,00061 | + | | | 73 |
| *Plekhh1* | <0.0001 | | + | | 74 |
| *Plp1* | <0.0001 | + | + | | 111,131,132 |
| *Rasd2* | <0.0001 | + | + | + | 133,134 |
| *Rtel1* | 0,00073 | + | | | 73 |
| *Tac1* | 0,00072 | + | | | 73 |
| *Th* | <0.0001 | | + | | 135-137 |
| *Thpo* | 0,0005 | + | | | 73 |
| *Trf* | <0.0001 | + | + | | 84,111,113,138-140 |
| *Trh* | <0.0001 | | + | | 74,141 |
| *Tspan2* | 0,00014 | + | | | 73 |
| *Xaf1* | <0.0001 | + | | | 73 |
