## Extended Data Table 2 for "Gene Knock Up via 3’UTR editing to study gene function *in vivo*"

Mätlik *et al.* Gene Knock Up via 3'UTR editing to study gene function *in vivo*

List of studies associating *GDNF* or GDNF receptor *GFRA1* (GDNF family receptor alpha, 1) gene or locus variants with schizophrenia.

| *Observation* | *Reference* |
|---|---|
| Suggestive evidence of linkage at 5p14.3-q11.2 in European ancestry families # | 142 |
| Suggestive evidence of linkage at 5p13 in a Costa Rican sample # | 143 |
| Genetic linkage at 5p13.3-13.2 within the interval between markers D5S1993 and D5S631 in a Puerto Rican family # | 144 |
| Association of a $(AGG)_n$ repeat polymorphism in the 3'UTR of *GDNF* gene | 145 |
| No significant association of 9 genotyped SNPs and $(AGG)_n$ repeat polymorphism in the *GDNF* gene | 146 |
| No significant association of $(AGG)_n$ repeat polymorphism in the *GDNF* gene in a Japanese sample | 147 |
| Association of GDNF family receptor alpha 1 (*GFRA1*) rs11197557 variant with schizophrenia | 148 |

### *GDNF* gene is located at 5p13.2
