## Extended Data Table 3 for "Gene Knock Up via 3’UTR editing to study gene function *in vivo*"

Mätlik *et al.* Gene Knock Up via 3'UTR editing to study gene function *in vivo*

| | **gRNA target sequences** |
|---|---|
| **MmGdnf gRNA A** | CCTGCTACAGTGCGAAGAAA |
| **MmGdnf gRNA B** | CAGTGCGAAGAAAGGGACCA |
| **MmGdnf gRNA C** | CATCTCGAGCAGGTTCGAAT |
| **MmGdnf gRNA D** | CAGCAGGGTAAAGTTTGCGA |
| **HsGdnf gRNA 1** | CCTGCTACAGTGCAAAGAAA |
| **HsGdnf gRNA 2** | CAGTGCAAAGAAAGGGACCA |
| **HsGdnf gRNA 3** | GCACTCCTTACGGTATCAGA |
| **HsGdnf gRNA 4** | ATAAAGTCTGTAGAGAACTT |

| | **Primer sequences for genotyping** |
|---|---|
| *Gdnf* $^{cKU}$ allele | |
| **F1** | TCTAAGAAAGCATTCCGCTAAACG |
| **F2** | TTCCAGGGTCAAGGAAGGCAC |
| **R1** | GGATGCGGTGGGCTCTATG |
| **R2** | TCCGCCATCTTGGTCCTTATC |
| *Gdnf* $^{KU}$ allele | |
| **F1** | GTGAATCGGCCGAGACAATG |
| **F2** | AGATGTCGTTCCAGACCCTCT |
| **R** | TGATGTCTCGGTCGTTTGTGTTA |

| | **qPCR primer sequences** |
|---|---|
| Mm Actb F | CTAAGGCCAACCGTGAAAAG |
| Mm Actb R | ACCAGAGGCATACAGGGACA |
| Mm Gdnf F | CGCTGACCAGTGACTCCAATATGC |
| Mm Gdnf R | TGCCGCTTGTTTATCTGGTGACC |
| Mm Drd1a F | GCGTGGTCTCCCAGATCG |
| Mm Drd1a R | GCATTTCTCCTTCAAGCCCCT |
| Mm Drd2 F | ACACACGCTACAGCTCCAAG |
| Mm Drd2 R | GGAGTAGACCACGAAGGCAG |
| Mm Gapdh F | GCCTCGTCCCGTAGACAAAA |
| Mm Gapdh R | ATGAAGGGGTCGTTGATGGC |
| Mm Hprt1 F | CAGTCCCAGCGTCGTGATTA |
| Mm Hprt1 R | TGGCCTCCCATCTCCTTCAT |
| Mm Pgk1 F | TTGGACAAGCTGGACGTGAA |
| Mm Pgk1 R | AACGGACTTGGCTCCATTGT |
| Mm Penk F | CCCAGGCGACATCAATTT |
| Mm Penk R | TCTCCCAGATTTTGAAAGAAGG |
| Mm Chat F | AAATGGCGTCCAACGAGGAT |
| Mm Chat R | GCTCGATCATGTCCAGGGAG |
| Mm Clic6 F | AATCCACCGAACACGAGGAG |
| Mm Clic6 R | ATGCTCTCGCCGTCATAACC |
| Hs Actb F | CCAACCGCGAGAAGATGA |
| Hs Actb R | CCAGAGGCGTACAGGGATAG |
| Hs Gdnf F | ATGTCCAACCTAGGGTCTGC |
| Hs Gdnf R | CATCCCATAACTTCATCTTAAAGTCC |
