## Extended Data Table 4 for "Gene Knock Up via 3’UTR editing to study gene function *in vivo*"

Mätlik *et al.* Gene Knock Up via 3'UTR editing to study gene function *in vivo*

### Main figures

| Panel | Groups | Group size | Sex | Data point description | Data exclusion | Statistical test: One way ANOVA | | Comments |
|---|---|---|---|---|---|---|---|---|
| | | | | | | ANOVA summary | Brown-Forsythe test | |
| 2a | wt/wt ±Cre | 6 | n.a | Each data point corresponds to a single animal. All samples were run in duplicates and averaged. | No data points were excluded from the analysis. | F=21.96 | F(DFn,DFd)=12.25 (2, 9) | Samples analysed in Fig. 2a-b originate from the same animals. |
| | wt/cKU -Cre | 2 | n.a | | | P=0.0003 | P=0.0027 | |
| | wt/cKU +Cre | 4 | n.a | | | R sq=0.8299 | | |

Post-hoc test: Tukey's multiple comparisons test

| Test details | Mean 1 | Mean 2 | Mean Diff, | SE of diff, | n1 | n2 | q | DF | Adj P Value |
|---|---|---|---|---|---|---|---|---|---|
| wt/wt ±Cre vs. wt/cKU -Cre | 1 | 1.235 | -0.2353 | 1.149 | 6 | 2 | 0.2895 | 9 | 0.9772 |
| wt/wt ±Cre vs. wt/cKU +Cre | 1 | 6.768 | -5.768 | 0.9085 | 6 | 4 | 8.978 | 9 | 0.0004 |
| wt/cKU -Cre vs. wt/cKU +Cre | 1.235 | 6.768 | -5.533 | 1.219 | 2 | 4 | 6.419 | 9 | 0.0036 |

| Panel | Groups | Group size | Sex | Data point description | Data exclusion | Statistical test: One way ANOVA | | Comments |
|---|---|---|---|---|---|---|---|---|
| | | | | | | ANOVA summary | Brown-Forsythe test | |
| 2b | wt/wt ±Cre | 4 | n.a | Each data point corresponds to a single animal. All samples were run in duplicates and averaged. | No data points were excluded from the analysis. | F=20.5 | F(DFn,DFd)=2.529 (2, 7) | Samples analysed in Fig. 2a-b originate from the same animals. |
| | wt/cKU -Cre | 2 | n.a | | | P=0.0012 | P=0.1490 | |
| | wt/cKU +Cre | 4 | n.a | | | R sq=0.8542 | | |

Post-hoc test: Tukey's multiple comparisons test

| Test details | Mean 1 | Mean 2 | Mean Diff, | SE of diff, | n1 | n2 | q | DF | Adj P Value |
|---|---|---|---|---|---|---|---|---|---|
| wt/wt ±Cre vs. wt/cKU -Cre | 147.2 | 149.5 | -2.314 | 74.48 | 4 | 2 | 0.0439 | 7 | 0.9995 |
| wt/wt ±Cre vs. wt/cKU +Cre | 147.2 | 503.4 | -356.2 | 60.81 | 4 | 4 | 8.285 | 7 | 0.0016 |
| wt/cKU -Cre vs. wt/cKU +Cre | 149.5 | 503.4 | -353.9 | 74.48 | 2 | 4 | 6.72 | 7 | 0.0051 |

| Panel | Groups | Group size | Sex | Data point description | Data exclusion | Statistical test: One way ANOVA | |
|---|---|---|---|---|---|---|---|
| 2e | wt/wt ±Cre | 8 | n.a | Each data point corresponds to a single animal. All samples were run in duplicates and averaged. | No data points were excluded from the analysis. | **ANOVA summary** | **Brown-Forsythe test** |
| | wt/cKU -Cre | 2 | n.a | | | F=19.8 | F(DFn,DFd)=9.123 (2, 13) |
| | wt/cKU +Cre | 6 | n.a | | | P=0.0001 | P=0.0033 |
| | | | | | | R sq=0.7529 | |

**Post-hoc test: Tukey's multiple comparisons test**

| Test details | Mean 1 | Mean 2 | Mean Diff, | SE of diff, | n1 | n2 | q | DF | Adj P Value |
|---|---|---|---|---|---|---|---|---|---|
| wt/wt ±Cre vs. wt/cKU -Cre | 1 | 1.017 | -0.01742 | 0.8561 | 8 | 2 | 0.0288 | 13 | 0.9998 |
| wt/wt ±Cre vs. wt/cKU +Cre | 1 | 4.523 | -3.523 | 0.5849 | 8 | 6 | 8.518 | 13 | 0.0001 |
| wt/cKU -Cre vs. wt/cKU +Cre | 1.017 | 4.523 | -3.505 | 0.8842 | 2 | 6 | 5.606 | 13 | 0.0043 |

| Panel | Groups | Group size | Sex | Data point description | Data exclusion | Statistical test: One way ANOVA | | Comments |
|---|---|---|---|---|---|---|---|---|
| 3a | wt | 17 | F (9), M (8) | Each data point corresponds to a single animal. All samples were run in duplicates and averaged. | No data points were excluded from the analysis. | **ANOVA summary** | **Brown-Forsythe test** | Wt group includes wt/wt ±Cre and wt/cKU -Cre genotypes. |
| | wt/cKU +Cre | 11 | F (5), M (6) | | | F=33.73 | F(DFn,DFd)=9.405 (2, 32) | |
| | cKU/cKU +Cre | 7 | F (3), M (4) | | | P<0.0001 | P=0.0006 | |
| | | | | | | R sq=0.6782 | | |

**Post-hoc test: Tukey's multiple comparisons test**

| Test details | Mean 1 | Mean 2 | Mean Diff, | SE of diff, | n1 | n2 | q | DF | Adj P Value |
|---|---|---|---|---|---|---|---|---|---|
| wt vs. wt/cKU +Cre | 1 | 1.985 | -0.9846 | 0.2412 | 17 | 11 | 5.772 | 32 | 0.0008 |
| wt vs. cKU/cKU +Cre | 1 | 3.265 | -2.265 | 0.28 | 17 | 7 | 11.44 | 32 | <0,0001 |
| wt/cKU +Cre vs. cKU/cKU +Cre | 1.985 | 3.265 | -1.28 | 0.3014 | 11 | 7 | 6.005 | 32 | 0.0005 |

| Panel | Groups | Group size | Sex | Data point description | Data exclusion | Statistical test: One way ANOVA | | Comments |
|---|---|---|---|---|---|---|---|---|
| 3c | wt | 10 | M | Each data point corresponds to a single animal. | No data points were excluded from the analysis. | ANOVA summary | Brown-Forsythe test | Wt group includes wt/wt ±Cre and wt/cKU -Cre genotypes. Samples analysed in Fig. 3c, 3d, 4a, 4b and Extended Data Fig. 2f originate from the same animals. |
| | wt/cKU +Cre | 7 | M | | | F=6.513 | F(DFn,DFd)=0.3064 (2, 18) | |
| | cKU/cKU +Cre | 4 | M | | | P=0.0074 | P=0.7398 | |
| | | | | | | R sq=0.4198 | | |

Post-hoc test: Tukey's multiple comparisons test

| Test details | Mean 1 | Mean 2 | Mean Diff, | SE of diff, | n1 | n2 | q | DF | Adj P Value |
|---|---|---|---|---|---|---|---|---|---|
| wt vs. wt/cKU +Cre | 12332 | 16756 | -4424 | 1622 | 10 | 7 | 3.856 | 18 | 0.0352 |
| wt vs. cKU/cKU +Cre | 12332 | 18478 | -6146 | 1947 | 10 | 4 | 4.463 | 18 | 0.0144 |
| wt/cKU +Cre vs. cKU/cKU +Cre | 16756 | 18478 | -1722 | 2063 | 7 | 4 | 1.181 | 18 | 0.6868 |

| Panel | Groups | Group size | Sex | Data point description | Data exclusion | Statistical test: One way ANOVA | | Comments |
|---|---|---|---|---|---|---|---|---|
| 3d | wt | 10 | M | Each data point corresponds to a single animal. | No data points were excluded from the analysis. | ANOVA summary | Brown-Forsythe test | Wt group includes wt/wt ±Cre and wt/cKU -Cre genotypes. Samples analysed in Fig. 3c, 3d, 4a, 4b and Extended Data Fig. 2f originate from the same animals. |
| | wt/cKU +Cre | 7 | M | | | F=3.551 | F(DFn,DFd)=0.3681 (2, 19) | |
| | cKU/cKU +Cre | 5 | M | | | P=0.0489 | P=0.6969 | |
| | | | | | | R sq=0.2721 | | |

Post-hoc test: Tukey's multiple comparisons test

| Test details | Mean 1 | Mean 2 | Mean Diff, | SE of diff, | n1 | n2 | q | DF | Adj P Value |
|---|---|---|---|---|---|---|---|---|---|
| wt vs. wt/cKU +Cre | 940 | 971.2 | -31.27 | 80.73 | 10 | 7 | 0.5478 | 19 | 0.921 |
| wt vs. cKU/cKU +Cre | 940 | 1173 | -232.6 | 89.73 | 10 | 5 | 3.666 | 19 | 0.045 |
| wt/cKU +Cre vs. cKU/cKU +Cre | 971.2 | 1173 | -201.4 | 95.92 | 7 | 5 | 2.969 | 19 | 0.1167 |

| Panel | Groups | Group size | Sex | Data point description | Data exclusion | Statistical test: Two way repeated measures ANOVA | | | | | | |
|---|---|---|---|---|---|---|---|---|---|---|---|---|
| 3e | wt | N=8, n=17 | Male | | | Source of variation | % of total variation | SS | DF | MS | F (DFn, DFd) | P value |
| | wt/cKU +Cre | N=8, n=20 | Male | Each data point corresponds to a measurement from single striatal slice. | Values obtained from one animal in wt group were excluded based on Grubbs test for outliers. | Interaction | 0.1817 | 0.04883 | 6 | 0.008139 | F (6, 132) = 4,082 | P=0,0009 |
| | cKU/cKU +Cre | N=4, n=10 | Male | | | Trial | 1.531 | 0.4114 | 3 | 0.1371 | F (3, 132) = 68,78 | P<0,0001 |
| | | | | | | Genotype | 15.54 | 4.177 | 2 | 2.088 | F (2, 44) = 4,173 | P=0,0219 |
| | | | | | | Subjects (matching) | 81.92 | 22.02 | 44 | 0.5004 | F (44, 132) = 251 | P<0,0001 |
| | | | | | | Residual | | 0.2632 | 132 | 0.001994 | | |

Post-hoc test: Tukey's multiple comparisons test, adjusted P-values

| Test details | Peak 1 | Peak 2 | Peak 3 | Peak 4 |
|---|---|---|---|---|
| wt vs. wt/cKU +Cre | 0.0542 | 0.0542 | 0.0723 | 0.0644 |
| wt vs. cKU/cKU +Cre | 0.0077 | 0.0284 | 0.0411 | 0.0648 |
| wt/cKU +Cre vs. cKU/cKU +Cre | 0.4902 | 0.7794 | 0.8037 | 0.9166 |

**Comments**

Data in Fig. 3e and 3f originate from the same experiment.

| Panel | Groups | Group size | Sex | Data point description | Data exclusion | Statistical test: Two way repeated measures ANOVA | | | | | | |
|---|---|---|---|---|---|---|---|---|---|---|---|---|
| 3f | wt | N=8, n=17 | Male | | | Source of variation | % of total variation | SS | DF | MS | F (DFn, DFd) | P value |
| | wt/cKU +Cre | N=8, n=20 | Male | Each data point corresponds to a measurement from single striatal slice. | Values obtained from one animal in wt group were excluded based on Grubbs test for outliers. | Interaction | 0.03768 | 1.669E-08 | 6 | 2.78E-09 | F (6, 132) = 1,203 | P=0,3089 |
| | cKU/cKU +Cre | N=4, n=10 | Male | | | Trial | 0.4183 | 1.853E-07 | 3 | 6.18E-08 | F (3, 132) = 26,7 | P<0,0001 |
| | | | | | | Genotype | 13.44 | 0.000005953 | 2 | 2.98E-06 | F (2, 44) = 3,461 | P=0,0402 |
| | | | | | | Subjects (matching) | 85.44 | 0.00003784 | 44 | 8.6E-07 | F (44, 132) = 371,8 | P<0,0001 |
| | | | | | | Residual | | 3.053E-07 | 132 | 2.31E-09 | | |

Post-hoc test: Tukey's multiple comparisons test, adjusted P-values

| Test details | Peak 1 | Peak 2 | Peak 3 | Peak 4 |
|---|---|---|---|---|
| wt vs. wt/cKU +Cre | 0.0473 | 0.0469 | 0.0406 | 0.0445 |
| wt vs. cKU/cKU +Cre | 0.0752 | 0.1284 | 0.1168 | 0.1494 |
| wt/cKU +Cre vs. cKU/cKU +Cre | 0.9727 | 0.9995 | 0.9994 | 0.9914 |

**Comments**

Data in Fig. 3e and 3f originate from the same experiment.

| Panel | Groups | Group size | Sex | Data point description | Data exclusion | Statistical test: One way ANOVA | | Comments |
|---|---|---|---|---|---|---|---|---|
| 4a | wt | 10 | M | Each data point corresponds to a single animal. | No data points were excluded from the analysis. | ANOVA summary | Brown-Forsythe test | Wt group includes wt/wt ±Cre and wt/cKU -Cre genotypes. Samples analysed in Fig. 3c, 3d, 4a, 4b and Extended Data Fig. 2f originate from the same animals. |
| | wt/cKU +Cre | 7 | M | | | F=5.131 | F(DFn,DFd)=1.464 (2, 19) | |
| | cKU/cKU +Cre | 5 | M | | | P=0.0165 | P=0.2562 | |
| | | | | | | R sq=0.3507 | | |

Post-hoc test: Tukey's multiple comparisons test

| Test details | Mean 1 | Mean 2 | Mean Diff, | SE of diff, | n1 | n2 | q | DF | Adj P Value |
|---|---|---|---|---|---|---|---|---|---|
| wt vs. wt/cKU +Cre | 833.6 | 391.6 | 442 | 169 | 10 | 7 | 3.698 | 19 | 0.043 |
| wt vs. cKU/cKU +Cre | 833.6 | 328.7 | 504.9 | 187.9 | 10 | 5 | 3.801 | 19 | 0.0371 |
| wt/cKU +Cre vs. cKU/cKU +Cre | 391.6 | 328.7 | 62.92 | 200.8 | 7 | 5 | 0.443 | 19 | 0.9475 |

| Panel | Groups | Group size | Sex | Data point description | Data exclusion | Statistical test: One way ANOVA | | Comments |
|---|---|---|---|---|---|---|---|---|
| 4b | wt | 10 | M | Each data point corresponds to a single animal. All samples were run in duplicates and averaged. | No data points were excluded from the analysis. | ANOVA summary | Brown-Forsythe test | Wt group includes wt/wt ±Cre and wt/cKU -Cre genotypes. Samples analysed in Fig. 3c, 3d, 4a, 4b and Extended Data Fig. 2f originate from the same animals. |
| | wt/cKU +Cre | 6 | M | | | F=9.379 | F(DFn,DFd)=4.225 (2, 17) | |
| | cKU/cKU +Cre | 4 | M | | | P=0.0018 | P=0.0324 | |
| | | | | | | R sq=0.5246 | | |

Post-hoc test: Tukey's multiple comparisons test

| Test details | Mean 1 | Mean 2 | Mean Diff, | SE of diff, | n1 | n2 | q | DF | Adj P Value |
|---|---|---|---|---|---|---|---|---|---|
| wt vs. wt/cKU +Cre | 1 | 0.3411 | 0.6589 | 0.1727 | 10 | 6 | 5.396 | 17 | 0.0037 |
| wt vs. cKU/cKU +Cre | 1 | 0.3697 | 0.6303 | 0.1979 | 10 | 4 | 4.505 | 17 | 0.0142 |
| wt/cKU +Cre vs. cKU/cKU +Cre | 0.3411 | 0.3697 | -0.02867 | 0.2159 | 6 | 4 | 0.1878 | 17 | 0.9903 |

| Panel | Groups | Group size | Sex | Data point description | Data exclusion | Statistical test: Two way ANOVA | | | | | | |
|---|---|---|---|---|---|---|---|---|---|---|---|---|
| 4g | wt/wt +Cre | 6 | M | Each data point corresponds to a single animal. | No data points were excluded from the analysis. | Source of variation | % of total variation | SS (Type III) | DF | MS | F (DFn, DFd) | P value |
| | cKU/cKU +Cre | 7 | M | | | Interaction | 0.009397 | 1.437 | 3 | 0.479 | F (3, 44) = 0,002808 | 0.9998 |
| | | | | | | Stimulus intensity | 32.36 | 4948 | 3 | 1649 | F (3, 44) = 9,67 | <0,0001 |
| | | | | | | Genotype | 18.37 | 2810 | 1 | 2810 | F (1, 44) = 16,47 | 0.0002 |
| | | | | | | Residual | | 7504 | 44 | 170.5 | | |

Post-hoc test: Sidak's multiple comparisons test, adjusted P-values

| Test details | 68 dB | 72 dB | 76 dB | 80 dB |
|---|---|---|---|---|
| wt/wt +Cre vs cKU/cKU +Cre | 0.1894 | 0.1622 | 0.2043 | 0.1678 |

| Panel | Groups | Group size | Sex | Data point description | Data exclusion | Statistical test: One way ANOVA | |
|---|---|---|---|---|---|---|---|
| 5b | wt/wt | 7 | M | Each data point corresponds to a single animal. All samples were run in duplicates and averaged. | No data points were excluded from the analysis. | ANOVA summary | Brown-Forsythe test |
| | wt/cKU | 7 | M | | | F=27.75 | F(DFn,DFd)=2.173 (2, 19) |
| | cKU/cKU | 8 | M | | | P<0.0001 | P=0.1414 |
| | | | | | | R sq=0.745 | |

Post-hoc test: Tukey's multiple comparisons test

| Test details | Mean 1 | Mean 2 | Mean Diff, | SE of diff, | n1 | n2 | q | DF | Adj P Value |
|---|---|---|---|---|---|---|---|---|---|
| wt/wt vs wt/cKU | 1 | 1.811 | -0.8111 | 0.1755 | 7 | 7 | 6.536 | 19 | 0.0005 |
| wt/wt vs cKU/cKU | 1 | 2.256 | -1.256 | 0.1699 | 7 | 8 | 10.45 | 19 | <0,0001 |
| wt/cKU vs cKU/cKU | 1.811 | 2.256 | -0.4447 | 0.1699 | 7 | 8 | 3.701 | 19 | 0.0428 |

| Panel | Groups | Group size | Sex | Data point description | Data exclusion | Statistical test: Two-tailed unpaired t-test | | F-test to compare variances | |
|---|---|---|---|---|---|---|---|---|---|
| 5c | wt/wt | 7 | M | Each data point corresponds to a single animal. | No data points were excluded from the analysis. | P value | t, df | F, DFn, Dfd | P-value |
| | cKU/cKU | 8 | M | | | 0.0121 | t=2.911, df=13 | 2.733, 6, 7 | 0.2082 |

| Panel | Groups | Group size | Sex | Data point description | Data exclusion |
|---|---|---|---|---|---|
| **5d** | wt/wt | 10 | M | Each data point corresponds to a single animal. | No data points were excluded from the analysis. |
| | cKU/cKU | 13 | M | | |

**Statistical test: Two way ANOVA**

| Source of variation | % of total variation | SS (Type III) | DF | MS | F (DFn, DFd) | P value |
|---|---|---|---|---|---|---|
| Interaction | 0.1758 | 64.15 | 3 | 21.38 | F (3, 84) = 0,05482 | P=0,9830 |
| Stimulus intensity | 7.63 | 2784 | 3 | 928.1 | F (3, 84) = 2,379 | P=0,0754 |
| Genotype | 2.133 | 778.6 | 1 | 778.6 | F (1, 84) = 1,996 | P=0,1614 |
| Residual | | 32765 | 84 | 390.1 | | |

| Panel | Groups | Group size | Sex | Data point description | Data exclusion | Statistical test: One way ANOVA | | Comments |
|---|---|---|---|---|---|---|---|---|
| | | | | | | ANOVA summary | Brown-Forsythe test | |
| **6d** | wt | 13 | n.a | Each data point corresponds to a single animal. All samples were run in duplicates and averaged. | No data points were excluded from the analysis. | F=13.62 | F(DFn,DFd)=2.87 (2, 18) | Samples analysed in Fig. 6d-f originate from the same animals. |
| | wt/KU | 6 | n.a | | | P=0.0003 | P=0.0828 | |
| | KU/KU | 2 | n.a | | | R sq=0.6021 | | |

**Post-hoc test: Tukey's multiple comparisons test**

| Test details | Mean 1 | Mean 2 | Mean Diff, | SE of diff, | n1 | n2 | q | DF | Adj P Value |
|---|---|---|---|---|---|---|---|---|---|
| wt vs wt/KU | 1 | 2.01 | -1.01 | 0.4408 | 13 | 6 | 3.239 | 18 | 0.083 |
| wt vs KU/KU | 1 | 4.432 | -3.432 | 0.6784 | 13 | 2 | 7.155 | 18 | 0.0002 |
| wt/KU vs KU/KU | 2.01 | 4.432 | -2.422 | 0.7292 | 6 | 2 | 4.698 | 18 | 0.0101 |

| Panel | Groups | Group size | Sex | Data point description | Data exclusion | Statistical test: One way ANOVA | | Comments |
|---|---|---|---|---|---|---|---|---|
| | | | | | | ANOVA summary | Brown-Forsythe test | |
| 6e | wt | 8 | n.a | Each data point corresponds to a single animal. All samples were run in duplicates and averaged. | No data points were excluded from the analysis. | F=8.003 | F(DFn,DFd)=3.164 (2, 13) | Samples analysed in Fig. 6d-f originate from the same animals. |
| | wt/KU | 6 | n.a | | | P=0.0054 | P=0.0759 | |
| | KU/KU | 2 | n.a | | | R sq=0.5518 | | |

Post-hoc test: Tukey's multiple comparisons test

| Test details | Mean 1 | Mean 2 | Mean Diff, | SE of diff, | n1 | n2 | q | DF | Adj P Value |
|---|---|---|---|---|---|---|---|---|---|
| wt vs wt/KU | 97.42 | 215.5 | -118 | 35.13 | 8 | 6 | 4.751 | 13 | 0.0132 |
| wt vs KU/KU | 97.42 | 254.9 | -157.4 | 51.43 | 8 | 2 | 4.329 | 13 | 0.0231 |
| wt/KU vs KU/KU | 215.5 | 254.9 | -39.41 | 53.12 | 6 | 2 | 1.049 | 13 | 0.7436 |

| Panel | Groups | Group size | Sex | Data point description | Data exclusion | Statistical test: Kruskal-Wallis one-way ANOVA | Comments |
|---|---|---|---|---|---|---|---|
| 6f | wt | 3 | n.a | Each data point corresponds to a single animal. All samples were run in duplicates and averaged. | No data points were excluded from the analysis. | P=0.0143 | Samples analysed in Fig. 6d-f originate from the same animals. |
| | wt/KU | 3 | n.a | | | KW statistic=6.846 | |
| | KU/KU | 6 | n.a | | | | |

Post-hoc test: Dunn's multiple comparisons test

| Test details | Rank 1 | Rank 2 | Rank Diff, | n1 | n2 | Adj P Value |
|---|---|---|---|---|---|---|
| wt vs wt/KU | 2 | 6.667 | -4.667 | 3 | 3 | 0.3388 |
| wt vs KU/KU | 2 | 8.667 | -6.667 | 3 | 6 | 0.0268 |
| wt/KU vs KU/KU | 6.667 | 8.667 | -2 | 3 | 6 | >0,9999 |

| Panel | Groups | Group size | Sex | Data point description | Data exclusion | Statistical test: Two-tailed unpaired t-test | | F-test to compare variances | |
|---|---|---|---|---|---|---|---|---|---|
| | | | | | | P value | t, df | F, DFn, Dfd | P-value |
| 6g | wt/wt | 6 | n.a | Each data point corresponds to one well of HEK-293 cells. All samples were run in duplicates and averaged. | Clones with both wt and KU alleles were not included to the analysis. | 0.0094 | t=3.292, df=9 | 1.45, 4, 5 | 0.6832 |
| | KU/KU | 5 | n.a | | | | | | |

### Extended Data Figures

| Panel | Groups | Group size | Sex | Data point description | Data exclusion | Statistical test: One way ANOVA | |
|---|---|---|---|---|---|---|---|
| ED 2a | wt/wt ±Cre | 6 | M | Each data point corresponds to a single animal. All samples were run in duplicates and averaged. | No data points were excluded from the analysis. | **ANOVA summary** | **Brown-Forsythe test** |
| | wt/cKU -Cre | 6 | M | | | F=173.7 | F(DFn,DFd)=8.392 (2, 13) |
| | wt/cKU +Cre | 4 | M | | | P<0.0001 | P=0.0046 |
| | | | | | | R sq=0.9639 | |

**Post-hoc test: Tukey's multiple comparisons test**

| Test details | Mean 1 | Mean 2 | Mean Diff, | SE of diff, | n1 | n2 | q | DF | Adj P Value |
|---|---|---|---|---|---|---|---|---|---|
| wt/wt ±Cre vs. wt/cKU -Cre | 66.14 | 81.96 | -15.82 | 6.19 | 6 | 6 | 3.615 | 13 | 0.0582 |
| wt/wt ±Cre vs. wt/cKU +Cre | 66.14 | 188.3 | -122.2 | 6.92 | 6 | 4 | 24.97 | 13 | <0,0001 |
| wt/cKU -Cre vs. wt/cKU +Cre | 81.96 | 188.3 | -106.4 | 6.92 | 6 | 4 | 21.74 | 13 | <0,0001 |

| Panel | Groups | Group size | Sex | Data point description | Data exclusion | Statistical test: One way ANOVA | | Comments |
|---|---|---|---|---|---|---|---|---|
| ED 2f | wt | 10 | M | Each data point corresponds to a single animal. All samples were run in duplicates and averaged. | Data points were excluded from the following groups: Drd1a (1 wt), Clic6 (1 cKU/cKU), Penk (1 wt, 1 wt/cKU) due to large difference between duplicates (SD>1) or based on Grubbs test for outliers. | **ANOVA summary** | **Brown-Forsythe test** | Wt group includes wt/wt ±Cre and wt/cKU -Cre genotypes. Samples analysed in Fig. 3c, 3d, 4a, 4b and Extended Data Fig. 2f originate from the same animals. |
| | wt/cKU +Cre | 6 | M | | | Drd1a | | |
| | cKU/cKU +Cre | 4 | M | | | F=5.114 | F(DFn,DFd)=1.067 (2, 16) | |
| | | | | | | P=0.0192 | P=0.3674 | |
| | | | | | | R sq=0.39 | | |
| | | | | | | Chat | | |
| | | | | | | F=4.963 | F(DFn,DFd)=4.569 (2, 17) | |
| | | | | | | P=0.0201 | P=0.0258 | |
| | | | | | | R sq=0.3687 | | |
| | | | | | | Clic6 | | |
| | | | | | | F=6.879 | F(DFn,DFd)=5.561 (2, 16) | |
| | | | | | | P=0.0070 | P=0.0147 | |
| | | | | | | R sq=0.4623 | | |
| | | | | | | Penk | | |
| | | | | | | F=7.415 | F(DFn,DFd)=0.5385 (2, 15) | |
| | | | | | | P=0.0058 | P=0.5945 | |
| | | | | | | R sq=0.4972 | | |

| Test details | Mean 1 | Mean 2 | Mean Diff, | SE of diff, | n1 | n2 | q | DF | Adj P Value |
|---|---|---|---|---|---|---|---|---|---|
| Post-hoc test: Tukey's multiple comparisons test | | | | | | | | | |
| Drd1a | | | | | | | | | |
| wt vs. wt/cKU +Cre | 1 | 0.7195 | 0.2805 | 0.1081 | 9 | 6 | 3.67 | 16 | 0.0486 |
| wt vs. cKU/cKU +Cre | 1 | 0.672 | 0.328 | 0.1233 | 9 | 4 | 3.763 | 16 | 0.0428 |
| wt/cKU +Cre vs. cKU/cKU +Cre | 0.7195 | 0.672 | 0.04748 | 0.1324 | 6 | 4 | 0.5071 | 16 | 0.9319 |
| Chat | | | | | | | | | |
| wt vs. wt/cKU +Cre | 1 | 0.4581 | 0.5419 | 0.1879 | 10 | 6 | 4.078 | 17 | 0.0265 |
| wt vs. cKU/cKU +Cre | 1 | 0.5385 | 0.4615 | 0.2153 | 10 | 4 | 3.032 | 17 | 0.1106 |
| wt/cKU +Cre vs. cKU/cKU +Cre | 0.4581 | 0.5385 | -0.08035 | 0.2349 | 6 | 4 | 0.4837 | 17 | 0.9378 |
| Clic6 | | | | | | | | | |
| wt vs. wt/cKU +Cre | 1 | 0.5781 | 0.4219 | 0.1475 | 10 | 6 | 4.045 | 16 | 0.0289 |
| wt vs. cKU/cKU +Cre | 1 | 0.4173 | 0.5827 | 0.188 | 10 | 3 | 4.382 | 16 | 0.0179 |
| wt/cKU +Cre vs. cKU/cKU +Cre | 0.5781 | 0.4173 | 0.1608 | 0.202 | 6 | 3 | 1.126 | 16 | 0.7108 |
| Penk | | | | | | | | | |
| wt vs. wt/cKU +Cre | 1 | 0.5825 | 0.4175 | 0.1182 | 9 | 5 | 4.996 | 15 | 0.008 |
| wt vs. cKU/cKU +Cre | 1 | 0.6667 | 0.3333 | 0.1273 | 9 | 4 | 3.702 | 15 | 0.0481 |
| wt/cKU +Cre vs. cKU/cKU +Cre | 0.5825 | 0.6667 | -0.08424 | 0.1421 | 5 | 4 | 0.8382 | 15 | 0.8261 |

| Panel | Groups | Group size | Sex | Data point description | Data exclusion | Statistical test: Two-tailed unpaired t-test | | F-test to compare variances | | Comments |
|---|---|---|---|---|---|---|---|---|---|---|
| ED 3a | wt/wt | N=6, n=10 | M | Each data point corresponds to a measurement from single striatal slice. | No data points were excluded from the analysis. | P value | t, df | F, DFn, Dfd | P-value | Extended Data Fig. 3a, 3c and 3d originate from the same experiment. |
| | wt/cKU | N=6, n=11 | M | | | 0.0187 | t=2.572, df=19 | 1.013, 10, 9 | 0.9934 | |

| Panel | Groups | Group size | Sex | Data point description | Data exclusion | Statistical test: One way ANOVA | |
|---|---|---|---|---|---|---|---|
| ED 3b | wt/wt | 7 | M | Each data point corresponds to a single animal. | No data points were excluded from the analysis. | ANOVA summary | Brown-Forsythe test |
| | wt/cKU | 7 | M | | | F=2.598 | F(DFn,DFd)=1.213 (2, 19) |
| | cKU/cKU | 8 | M | | | P=0.1006 | P=0.3193 |
| | | | | | | R sq=0.2147 | |

| Panel | Groups | Group size | Sex | Data point description | Data exclusion |
|---|---|---|---|---|---|
| ED 3c | wt/wt | N=6, n=10 | M | Each data point corresponds to a measurement from single striatal slice. | No data points were excluded from the analysis. |
| | wt/cKU | N=6, n=11 | M | | |

**Statistical test: Two way repeated measures ANOVA**

| Source of variation | % of total variation | SS | DF | MS | F (DFn, DFd) | P value | Comments |
|---|---|---|---|---|---|---|---|
| Interaction | 0.04372 | 0.01873 | 3 | 0.006244 | F (3, 57) = 0,5402 | P=0,6567 | Extended Data Fig. 3a, 3c and 3d originate from the same experiment. |
| Trial | 0.4872 | 0.2087 | 3 | 0.06957 | F (3, 57) = 6,019 | P=0,0012 | |
| Genotype | 0.02147 | 0.009197 | 1 | 0.009197 | F (1, 19) = 0,004166 | P=0,9492 | |
| Subjects (matching) | 97.9 | 41.94 | 19 | 2.208 | F (19, 57) = 191 | P<0,0001 | |
| Residual | | 0.6588 | 57 | 0.01156 | | | |

| Panel | Groups | Group size | Sex | Data point description | Data exclusion |
|---|---|---|---|---|---|
| ED 3d | wt/wt | N=6, n=10 | M | Each data point corresponds to a measurement from single striatal slice. | No data points were excluded from the analysis. |
| | wt/cKU | N=6, n=11 | M | | |

**Statistical test: Two way repeated measures ANOVA**

| Source of variation | % of total variation | SS | DF | MS | F (DFn, DFd) | P value | Comments |
|---|---|---|---|---|---|---|---|
| Interaction | 0.1041 | 5.167E-07 | 3 | 1.722E-07 | F (3, 57) = 0,4351 | P=0,7287 | Extended Data Fig. 3a, 3c and 3d originate from the same experiment. |
| Trial | 0.1523 | 7.562E-07 | 3 | 2.521E-07 | F (3, 57) = 0,6367 | P=0,5945 | |
| Genotype | 1.537 | 0.000007632 | 1 | 0.000007632 | F (1, 19) = 0,3118 | P=0,5831 | |
| Subjects (matching) | 93.66 | 0.0004651 | 19 | 0.00002448 | F (19, 57) = 61,83 | P<0,0001 | |
| Residual | | 0.00002257 | 57 | 3.959E-07 | | | |

| Panel | Groups | Group size | Sex | Data point description | Data exclusion | Statistical test: Two-tailed unpaired t-test | |
|---|---|---|---|---|---|---|---|
| ED 3e | wt/wt | 11 | M | Each data point corresponds to a single animal. | No data points were excluded from the analysis. | **P value** | **t, df** |
| | wt/Gdnfh | 8 | M | | | 0.4667 | t=0.7447, df=17.00 |
| | wt/wt | 8 | M | Each data point corresponds to a single animal. | No data points were excluded from the analysis. | **P value** | **t, df** |
| | MEN2B/MEN2B | 9 | M | | | 0.8986 | t=0.1296, df=15.00 |
